# cGMP signaling: probing antagonistic cyclic nucleotide platelet signals by modeling and experiment

**DOI:** 10.1101/2021.02.01.429103

**Authors:** Tim Breitenbach, Nils Englert, Özge Osmanoglu, Natalia Rukoyatkina, Gaby Wangorsch, Andreas Friebe, Elke Butt, Robert Feil, Marcus Dittrich, Stepan Gambaryan, Thomas Dandekar

**Author notes:** Email addresses: TB. ÖO (Özge Osmanoglu).

## Abstract

**Background:** The cyclic nucleotides cAMP and cGMP inhibit platelet activation.

**Results:** We extended an older model and systematically integrated drugs as external stimuli. Data driven modeling allowed us to design models that provide a quantitative output for quantitative input information. This relies on condensed information about involved regulation and modeling of pharmacological interventions by systematic optimization methods. By multi-experiment fitting, we validated our model optimizing the parameters of the model. In addition, we show how the output of the developed cGMP model can be used as input for a modular model of VASP phosphorylation and for the activity of cAMP and cGMP pathways in platelets.

**Conclusions:** We present a model for cGMP signaling and VASP phosphorylation, that allows to estimate drug action on any of the inhibitory cyclic nucleotide pathways (cGMP, cAMP) and has been validated by experimental data.

## Introduction

Platelets play a fundamental role in normal and pathological hemostasis. The fine-tuning between activating and inhibitory factors prevents spontaneous platelet activation, aggregation as well as occlusion of blood vessels. However, in the case of vascular injury, platelets adhere to injured endothelium and subendothelial matrix to form localized thrombi and prevent blood loss. A variety of factors, including collagen, thrombin, ADP, von Willebrand factor (vWF), thromboxane, podoplanin, and others, act by activation of corresponding receptors and utilizing multiple intracellular transduction mechanisms that promote platelet activation. On the other hand, endothelial cell derived prostacyclin and nitric oxide (NO) act by increasing platelet cAMP and cGMP concentrations, respectively, which mediate the major intracellular platelet inhibitory mechanisms. Elevated cAMP and cGMP levels activate corresponding protein kinases, protein kinase A (PKA) and protein kinase G (PKG), respectively, which phosphorylate multiple substrates responsible for platelet inhibition. One of the well-known substrates of both PKA and PKG in platelets is the vasodilator-stimulated phosphoprotein (VASP). Due to the commercial availability of phospho-specific antibodies against phosphorylated VASP, measurement of phospho-VASP by Western blot analysis has become a standard method to monitor the biochemical activity of the cAMP and cGMP pathways in platelets. The mechanisms of platelet inhibition via cAMP-PKA and cGMP-PKG signaling have been described in several reviews (Walter and Gambaryan 2009; Smolenski 2012; Gambaryan, Tsikas, 2015; Makhoul et al, 2018)Wen et al., 2017).

In an initial work by Wangorsch *et al*. (2011) we primarily analysed the action of cAMP on platelet function using ordinary differential equations (ODEs) to model cyclic nucleotide signaling. In that work, we focussed on the cAMP-pathway and the effect of different drugs on the cAMP levels. Our new study here focusses on the cGMP and extends our previous work by parametrizing the model according to experimental data including this specific cGMP pathways. Furthermore, we systematically included drugs as external stimuli by modeling and predicting the effects with mathematical functions.

The signaling pathway of cGMP in platelets is more complex than cAMP signaling as the molecule can activate PKG, stimulate or inhibit several phosphodiesterases (PDEs) where the ratio of active and inactive forms changes or activate cGMP-gated cation channels (although not present in platelets). Binding of NO to soluble guanylyl cyclase (sGC), leads to an increase in cGMP production that can be orders of magnitudes higher than in the unstimulated situation. sGC converts GTP into the second messenger cGMP, which binds to effectors such as PKG, phosphodiesterases (PDE), or cGMP-gated cation channels. The effects of an active NO/cGMP pathway in platelets include the inhibition of platelet aggregation and thrombosis. Recent data suggest that NO-induced cGMP generation in platelets is strongly potentiated by shear stress and that this mechanosensitive NO/cGMP/PKG cascade acts as an auto-regulatory brake of thrombosis (Wen et al., 2018). Pharmacologically, sGC can be stimulated in different ways: NO-releasing compounds like sodium nitroprusside (SNP) activate sGC in the same way as physiological NO released by endothelil cells. sGC stimulators such as BAY 41-2722 (Bay) and riociguat (BAY 63-2521; Rio) stimulate sGC independently of NO on the one hand and synergistically with NO on the other hand by stabilizing the activating conformational change. For NO stimulation, sGC must carry a reduced (Fe^2+^) prosthetic heme group. In contrast, synthetic sGC activators are able to stimulate “oxidized” sGC carrying an oxidized heme group (Fe^3+^) or even heme-free sGC. An overview of the NO/cGMP pathway in platelets is given in Figure 1.

**Figure 1.**
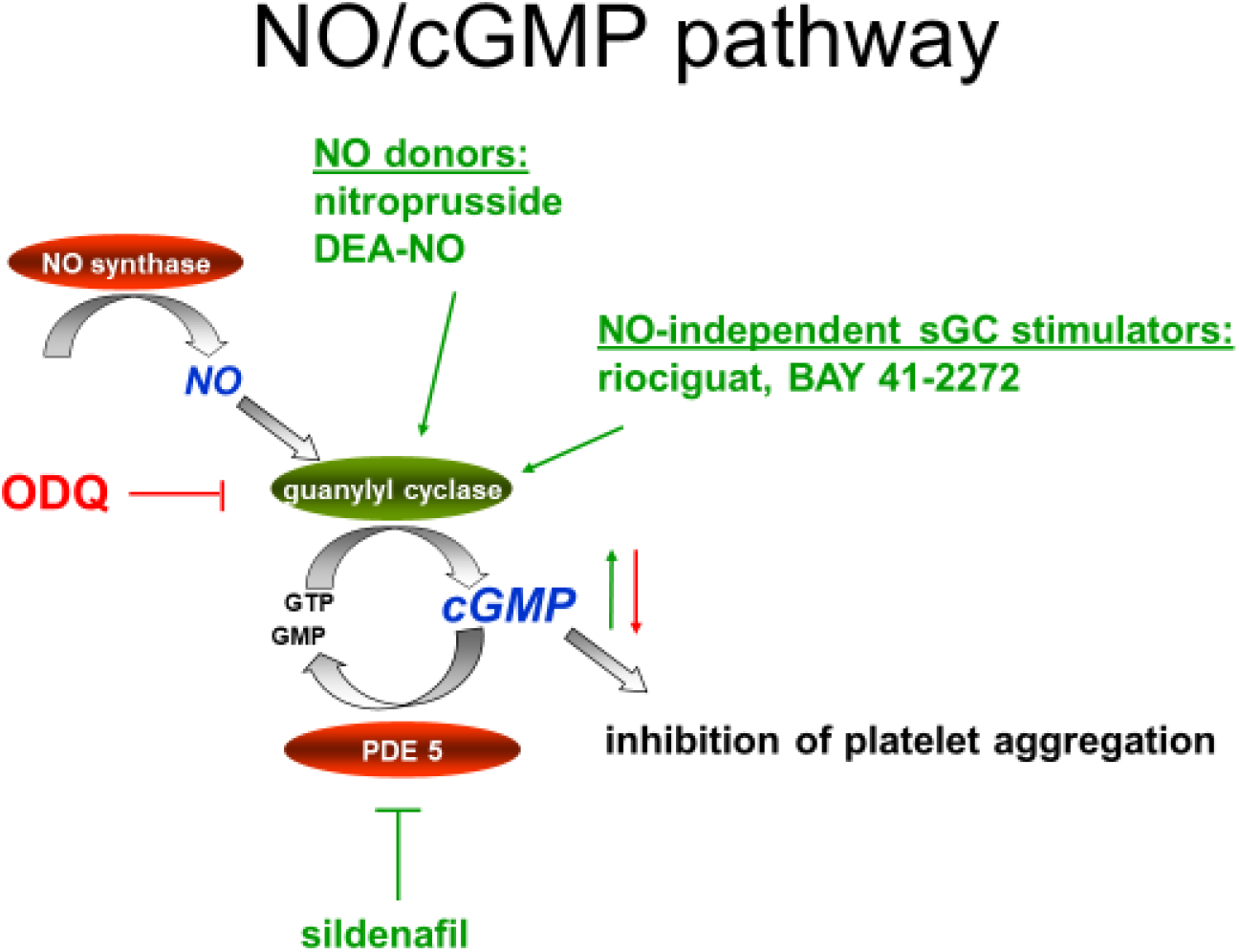
NO/cGMP signaling in platelets. NO is generated from L-arginine by NO synthases, for instance, in endothelial cells of the vessel wall. Then, NO diffuses into platelets and activates cGMP synthesis by soluble guanylyl cyclase. This enzyme can be pharmacologically inhibited by ODQ and stimulated by NO-releasing compounds or NO-independent stimulators. Increased cGMP inhibits platelet aggregation. The signaling cascade is terminated by cGMP degradation via PDE5. Inhibition of PDE5 by sildenafil results in elevated cGMP levels.

Recently, cGMP signaling in platelets was modeled by Kleppe et al. (2018). The authors discussed different variants of a core model, whereas we use a comprehensive, unifying approach for our study. Moreover, Kleppe et al (2018) present a purely theoretical model, while our model is directly validated and parametrized by experimental data.

By including cAMP signaling and external cGC stimuli, we present here a rational overarching framework to model cyclic nucleotide based platelet dynamics and to describe biological processes. In order to validate our framework, we have measured data of cAMP and cGMP levels in platelets - in particular cGMP concentration with respect to different sGC stimulatory drugs and drug concentrations.

We would like to model this relation in order to be able to calculate the drug exposure for desired cGMP concentration. Our intention in this work is not to fit a mathematical model to the data in order to obtain certain reaction constants from the model but to obtain a model that describes the cGMP concentration, its changes and its modulation sufficiently well.

We present a model that has been validated by experimental data and includes cGMP pathways as well as VASP phosphorylation. Our modeling of the inhibitory pathways helps to better understand the modulation of the fragile balance of platelet function, especially because inhibitory pathways are important to prevent platelet aggregation and cardiovascular pathology such as arterial thrombosis and cardiac infarction. Moreover, we present here a model which includes besides cGMP pathways also VASP phosphorylation and has been validated by experimental data. A further application of our model is that the cGMP levels can be calculated by the VASP levels. Once the VASP levels are given the optimization framework in *Breitenbach et al. (2019a,b)* can be used to calculate the cGMP level such that the predicted VASP level fits optimally to the given VASP levels. In this optimization scenario, the cGMP levels are defined as external stimuli that cause a corresponding VASP level.

## Results

### Setting up the model

To model platelet responses regarding the cGC-cGMP pathway, we extended the core model proposed in Wangorsch et al. (2011) (see section therein 2.4 “Differential equations of the dynamic variables”) and demonstrate how this mathematical model can be extended systematically to model the influence of external stimuli to cGMP pathways (Table 1).

**Table 1:**
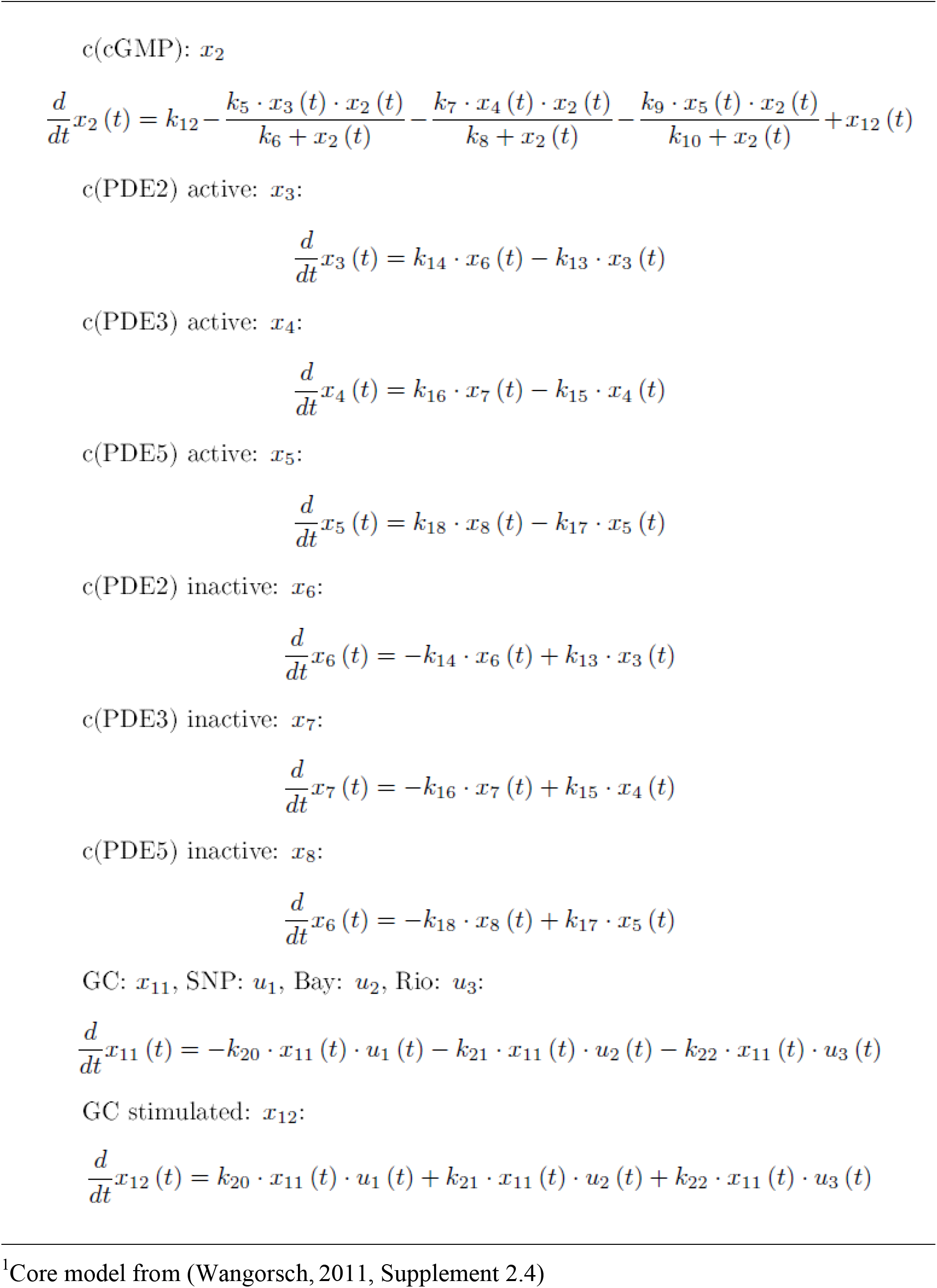
Our basic model for describing the cGMP level^1^.

In our case, these external stimuli are drugs, sodium nitroprussid (SNP), BAY 41-2272 (Bay) and riociguat (Rio), that all stimulate sGC. We adopt the same notations for the dynamic variables and the parameters in order to easily combine our extension into the already validated core model. Our aim is to show how to include our systematic approach for modeling the effect of drugs and pharmacologically active agents as external stimuli. For this purpose, we just focus on that part of the model that describes the dynamics of the cGMP production and generated data sets for drugs that interact with the sGC. Moreover, we combine module-wise the core model with models describing the phosphorylation of VASP. The notion of this procedure is to model parts of a biological system each as a function that maps input information to output information, to improve and to validate each sub model and thus systematically to assemble the total system from the subsystems connecting them by the input-output-interfaces.

Furthermore, as in the model and topology from Wangorsch et al. (2011) and confirmed by the experimental data and implemented in our model (see Table 2), there is no feedback from the equations of AMP and cAMP back into the equations for cGMP. Analogously, the value for the AMP concentration has according to the data no feedback into the network with regard to the cGMP production. In order to focus just on the equations necessary for our model of cGMP, we leave the equation for cAMP, AMP and GMP out of our discussion. Focusing on the necessary equations keeps the discussion clear for the intended purpose to demonstrate how to incorporate the drugs as external stimuli into modeling. Since the details about modeling the cGMP and cAMP levels in platelets with a corresponding system of equations for the dynamic variables can be found in Wangorsch *et al*. (2011), we started from the equations given by Wangorsch *et al*. (2011).

**Table 2:**
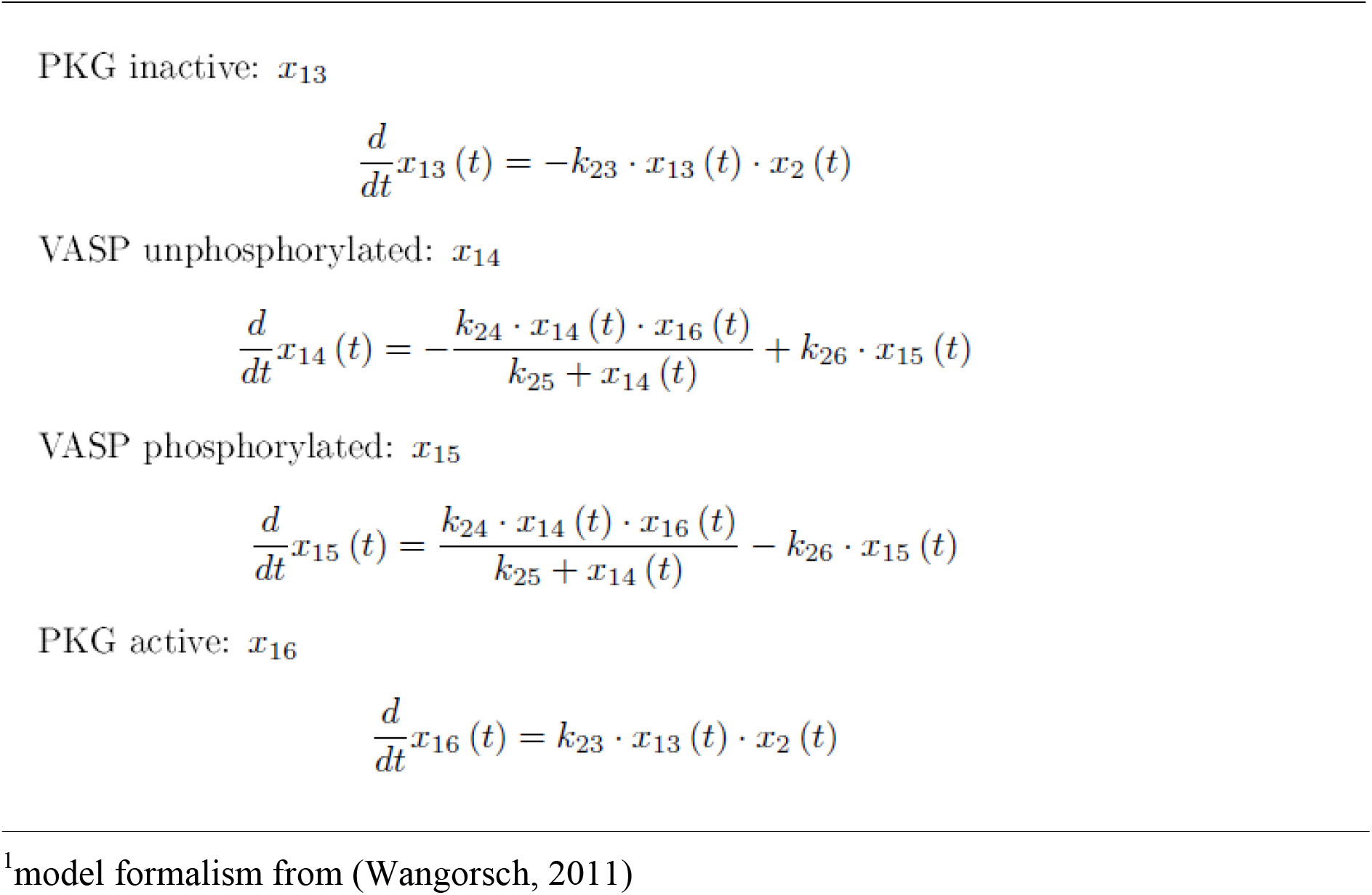
VASP phosphorylation^1^.

The models explained in Wangorsch et al. (2011) as well as analogous models can be extended with our presented method. We show our model in Table 2 and explain the modifications done to the model of Wangorsch et al. (2011) in the following:

The basic idea is that the drugs stimulate the GC such that the turnover of cGMP is increased. That implies that by the action of the drugs, the population of unbound GC is decreased and is turned into a species called GC-stimulated with an increased turnover of cGMP.

For this purpose, we hypothesize a mechanism of law of mass action assuming one molecule of drug binds to one molecule GC. In our model, we see that the functions that model SNP, *u*_1_, Bay, *u*_2_ and Rio, *u*_3_ are time dependent. In our experiments, we assume the concentration of drug to be constant. However, our model is not restricted to that assumption. In case of experimental capabilities to vary the concentration of the drugs time dependently, we can model this by the time dependent control functions *u*_1_, *u*_2_ and *u*_3_. If a drug itself is subject to a dynamic change, which is naturally the case in a patient, then the model can be extended as discussed in the Discussion.

The turnover of cGMP reaches a maximum when all the initial GC molecules are activated and turned into the GC-stimulated form. This means that the population of non-stimulated GC is small and the population of stimulated GC is big/large.

In our model, *k*_12_ models the basal GC afflux of cGMP. The increasing afflux of cGMP by CG stimulated is modeled by a further term *x*_12_ in the equation that describes the dynamic of cGMP (*x*_2_). Usually one would expect that the last two equations (for *x*_11_ and *x*_12_) model concentrations. Since we do not have any data for the concentrations of GC and GC-stimulated, we have the parameters *k*_20_, *k*_21_, *k*_22_ fit by Potterswheel such that the dynamic variables *x*_11_ and *x*_12_ correspond to the additional afflux. Once sufficient data become available to model also the concentration for GC and GC-stimulated, we’ll have to include another parameter *k*_23_ and add *k*_23_ ⋅ *x*_12_ instead of *x*_12_ in the equation for cGMP. In our case, we can omit this parameter *k*_23_ since we have no experimental data for *x*_12_ and thus build a model with less parameters. This is useful because a model with less parameters is considered generally as the better one (model reduction) providing it fits similar well to the experimental data.

In Table 2, we present a model that we use to describe the VASP phosphorylation. We refined the model of the supplement S1 Section 5 of Wangorsch *et al*. (2011). In the new model we use the concentration of cGMP, *x*_2_, as input for the model of Table 2.

In Table 3 we present a basal model (less free parameters than the model from Table 2) that we also fit to the data and thus show that also this basal model based on the law of mass action consisting of two terms is sufficient. We have a term that models the phosphorylation of VASP being proportional to the concentration of cGMP, *x*_2_ and the unphosphorylated VASP, *x*_14_. In addition, we have a term modeling the decay of the phosphorylated VASP, i.e. the dephosphorylation of the phosphorylated VASP, *x*_15_.

**Table 3:**
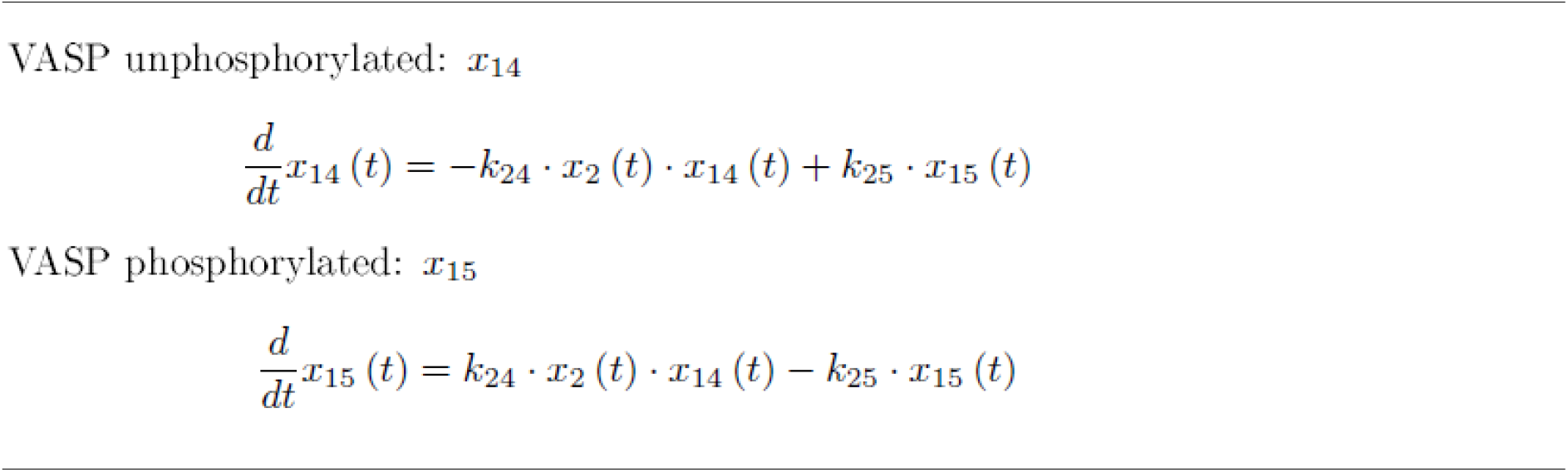
Basal model for the phosphorylation of VASP.

### Experimental investigation: cGMP time series data

An independent way to understand cGMP signaling are concentration- and time-dependent data from cGMP stimulation experiments. These experiments provide a general look into the changes in cGMP levels in response to different concentrations and incubation times of sGC agonists. They provide reliable information on activation levels of specific proteins in specific conditions (here VASP and PDE5 phosphorylation). Therefore, these experiments are very useful when the scope is narrowed down to individual proteins. However, if there are multiple conditions to test, in silico models are useful to narrow down the candidates. For example, here we want to test a couple of sGC agonist and inhibitors of downstream proteins.

In silico simulations help to find out which combination of these inputs may give the optimal results. The in silico findings can then be also experimentally validated. In these cases, in silico models and their simulation help narrow down the conditions that would require time- and labor-intensive experiments.

We first looked at the effect of sGC inhibition by ODQ on cGMP levels and downstream VASP phosphorylation (Figure 5). In the absence of sGC activator (SNP) and PDE5 inhibitor (Sildenafil), neither increase of cGMP/cAMP levels nor phosphorylation of VASP or PDE5 is observed.In the absence of ODQ, inhibition of PDE5 by Sildenafil showed very high levels of cGMP and VASP phosphorylation, pointing to higher PKG activity, while no change was observed on cAMP levels. Combination of ODQ and Sildenafil led to lower cGMP levels, PKG activity and VASP phosphorylation. Stimulation with SNP itself resulted in high cGMP levels and concomitant phosphorylation of PDE5 and VASP. However, in the presence of ODQ the effect of SNP is completely abolished.

**Figure 2.**
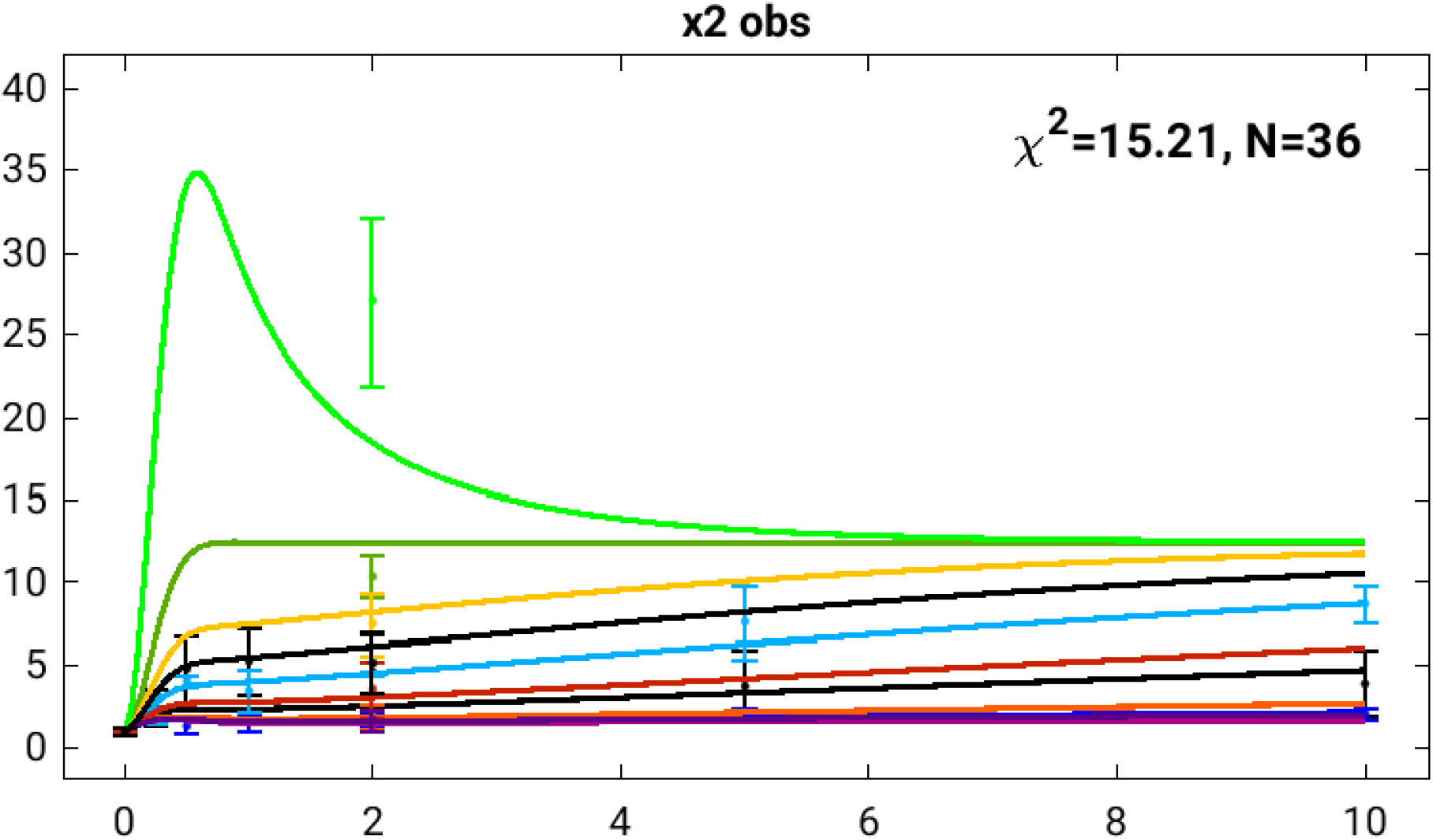
Time curves from the model with the best fitted parameters for different drugs (Rio, Bay, SNP) at different concentrations. The figure is a screenshot from Potterswheel and intended as an illustration of the resulting model parameters depicted in Table 4 with the data in the file data_cGMP.xls. On the x-axis there is the time in minutes and on the y-axis there is the cGMP fold increase with regard to the cGMP turnover of the GC without any administration of drug. Each colour belongs to one scenario with corresponding activated external stimuli, data points and error bars. In the presented case, the corresponding external stimuli are the presence of Bay, SNP and Rio in different concentrations. In each scenario the curve is the cGMP fold increase predicted by the model with the best fit of parameters. The corresponding data points are the mean values of all values at the specific point of time of any experiments done with the specific drug used in the scenario and the concentration of the drug. The error bar of each data point is the corresponding standard deviation. The time curves are as follows from top to bottom: (light green, Rio: 20 µM), (dark green, Rio: 5 µM), (yellow, Rio: 2.5 µM), (black, SNP: 5 µM), (light blue, Rio: 1 µM), (red, Rio: 0.5 µM), (black, SNP: 1 µM), (orange, Rio: 0.1 µM), (dark blue, Bay: 5 µM) (violet, SNP: 0.1), (pink, SNP: 0.01 µM). The **χ**^**2**^ - value is fine and shows a good fit (see Materials and Methods for details) The **χ**^**2**^ -value is a measure for the distance between the model prediction and the corresponding data points.

**Figure 3.**
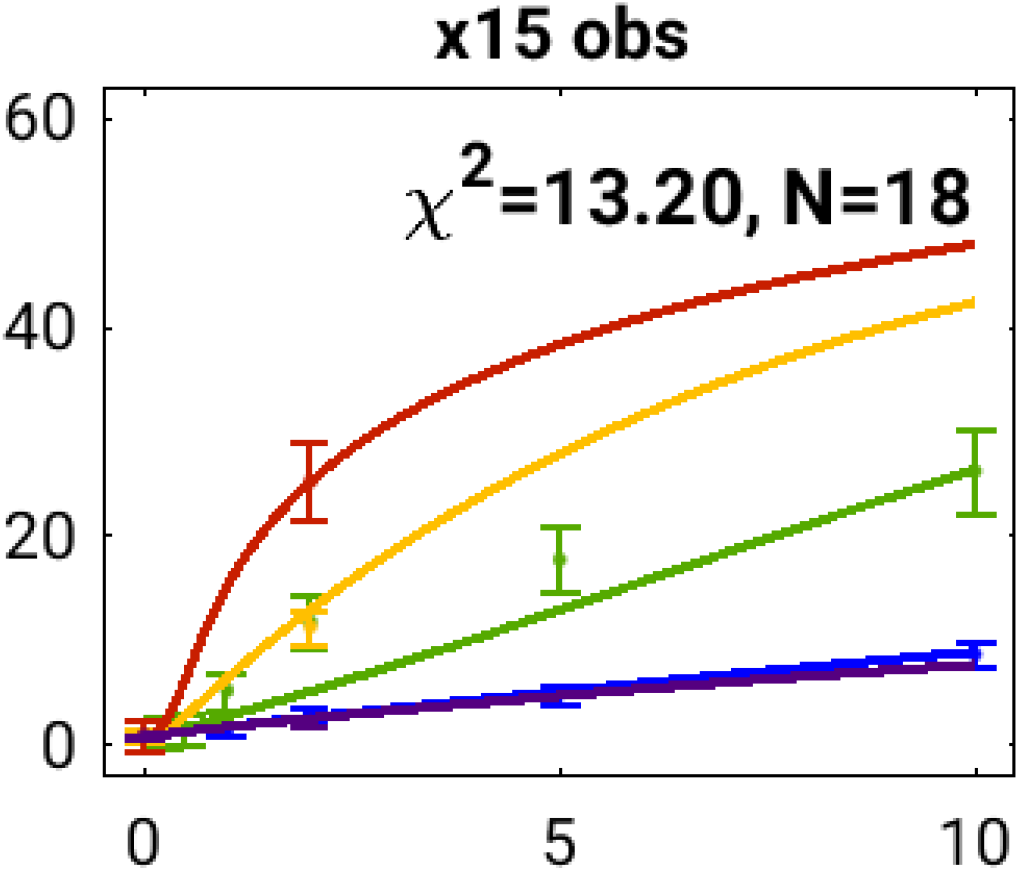
Time curves from the basal VASP model as depicted in Table 3. The figure is a screenshot of Potterswheel for the best fitting parameters for different drugs and intended as an illustration. The data in detail is found in the file data_cGMP_VASP.xls. On the x-axis we have the time in minutes and on the y-axis the ratio p-VASP/Actin. Each colour belongs to one scenario with corresponding activated external stimuli, data points and error bars. In the presented case, the corresponding external stimuli are the presence of Bay, SNP and Rio in different concentrations. In each scenario the curve is the ratio p-VASP/Actin predicted by the model with the best fit of parameters. The corresponding data points are the mean values of all values at the specific point of time of any experiments done with the specific drug used in the scenario and the concentration of the drug. The error bar of each data point is calculated according to the standard error model of Potterswheel as explained in the paragraph starting on page 7. The time curves are as follows from top to bottom: (red, Rio: 20 µM), (yellow, Rio: 5 µM), (green, Rio: 1 µM), (blue, Bay: 5 µM), (violet, SNP: 0.01 µM) The **χ**^**2**^ -value is a measure for the distance between the model prediction and the corresponding data points. Consequently, the smaller the value is, the more likely it is that the model explains the data. Considering the number of data points (N=18), the p-values can be calculated giving the probability to obtain the value of **χ**^**2**^ assuming that the hypothesis that the model explains the data is true. The p-value gives an estimation if the model based on the data has to be rejected or can be reasonably used for further calculations.

**Figure 4.**
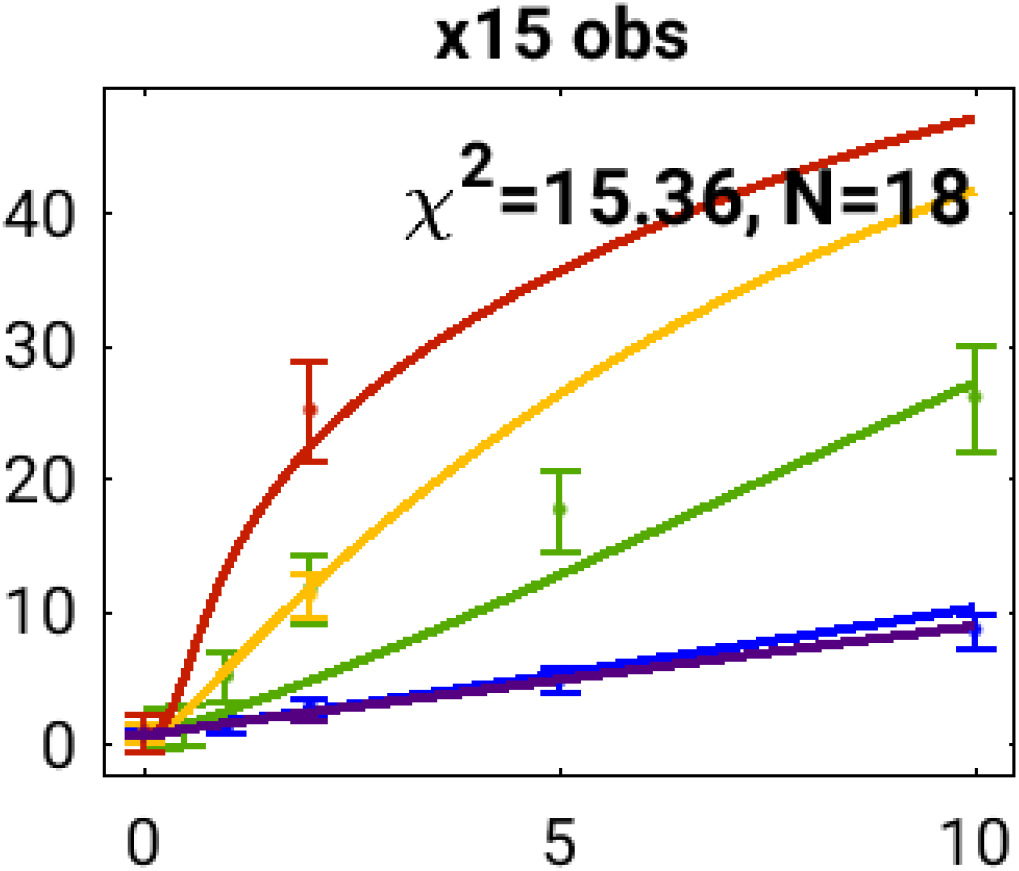
Time curves from the model as depicted in Table 2. The figure is a screenshot of Potterswheel for the best fitting parameters for different drugs and intended as an illustration. On the x-axis we have the time in minutes and on the y-axis the ratio p-VASP/Actin. Each colour belongs to one scenario with corresponding activated external stimuli, data points and error bars. In the presented case, the corresponding external stimuli are the presence of Bay, SNP and Rio in different concentrations. In each scenario the curve is the ratio p-VASP/Actin predicted by the model with the best fit of parameters. The corresponding data points are the mean values of all values at the specific point of time of any experiments done with the specific drug used in the scenario and the concentration of the drug. The error bar of each data point is calculated according to the standard error model of Potterswheel as explained in the paragraph starting on page 7. The time curves are as follows from top to bottom: (red, Rio: 20 µM), (yellow, Rio: 5 µM), (green, Rio: 1 µM), (blue, Bay: 5 µM), (violet, SNP: 0.01 µM) The **χ**^**2**^ -value a measure for the distance between the model prediction and the corresponding data points. Consequently, the smaller the value is, the more likely it is that the model explains the data. Considering the number of data points (N=18), the p-values can be calculated giving the probability to obtain the value of **χ**^**2**^ assuming that the hypothesis that the model explains the data is true. The p-value gives an estimation if the model based on the data has to be rejected or can be reasonably used for further calculations.

**Figure 5.**
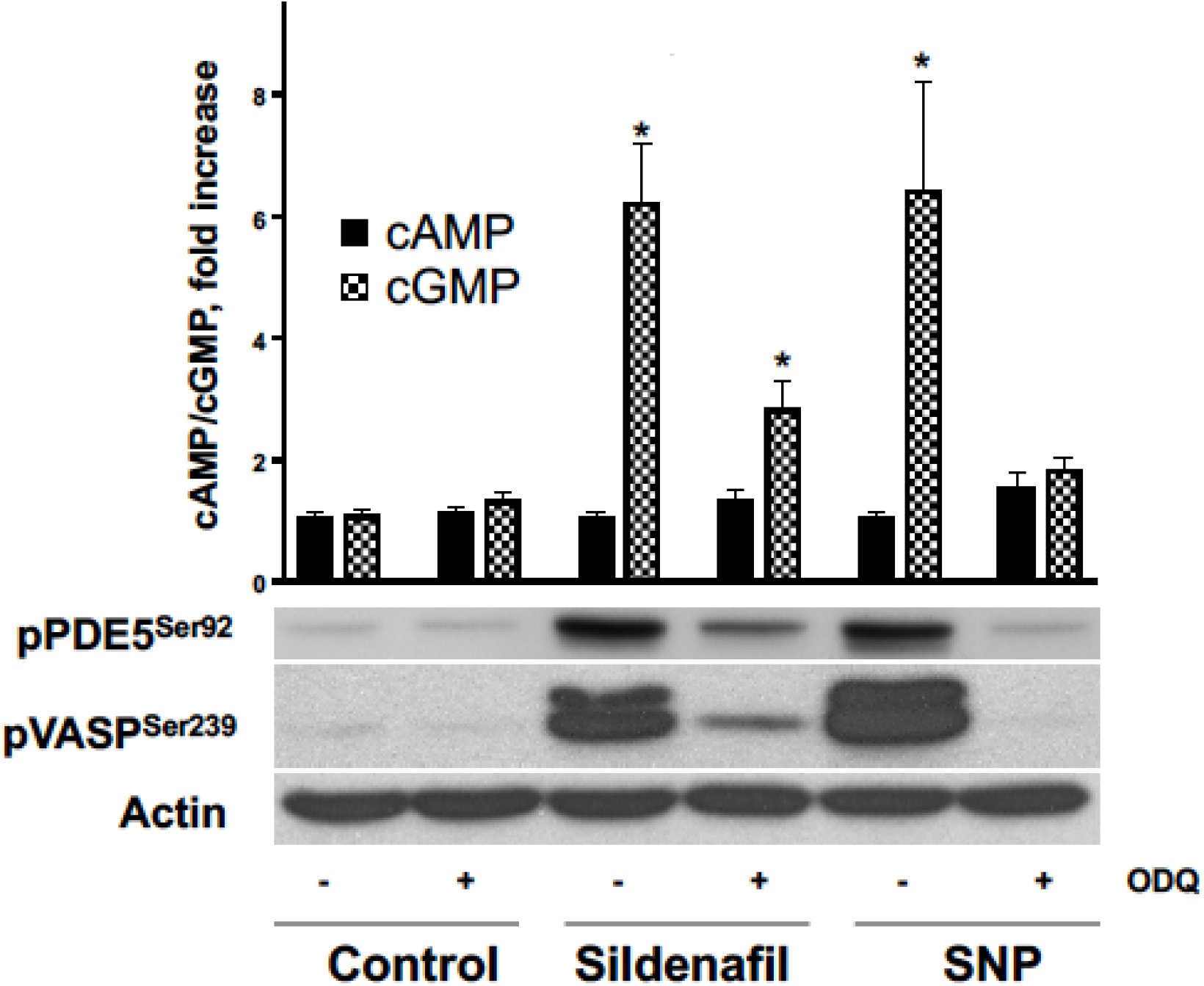
Inhibition of sGC by ODQ prevents cGMP accumulation and VASP phosphorylation in human platelets. Washed human platelets (3×10^8^/ml) were stimulated with ODQ (ODquinolaxin), Sildenafil (5 µM, 10 min) or SNP (5 µM, 1 min) with or without preincubation (ODQ 10 µM, 10 min). sGC activation was monitored by the analysis of PDE5 and VASP phosphorylation (pPDE5^Ser92^ and pVASP^Ser239^ Western blot, actin blot served as a loading control) and cAMP/ cGMP levels by EIA. Results of three independent experiments are presented as mean ± SEM. * p < 0.05 compared to control taken as 1. Shown is a representative blot from three independent experiments.

Next, we investigated the concentration-dependent effect of Riociguat on sGS activation (Figure 6).

**Figure 6.**
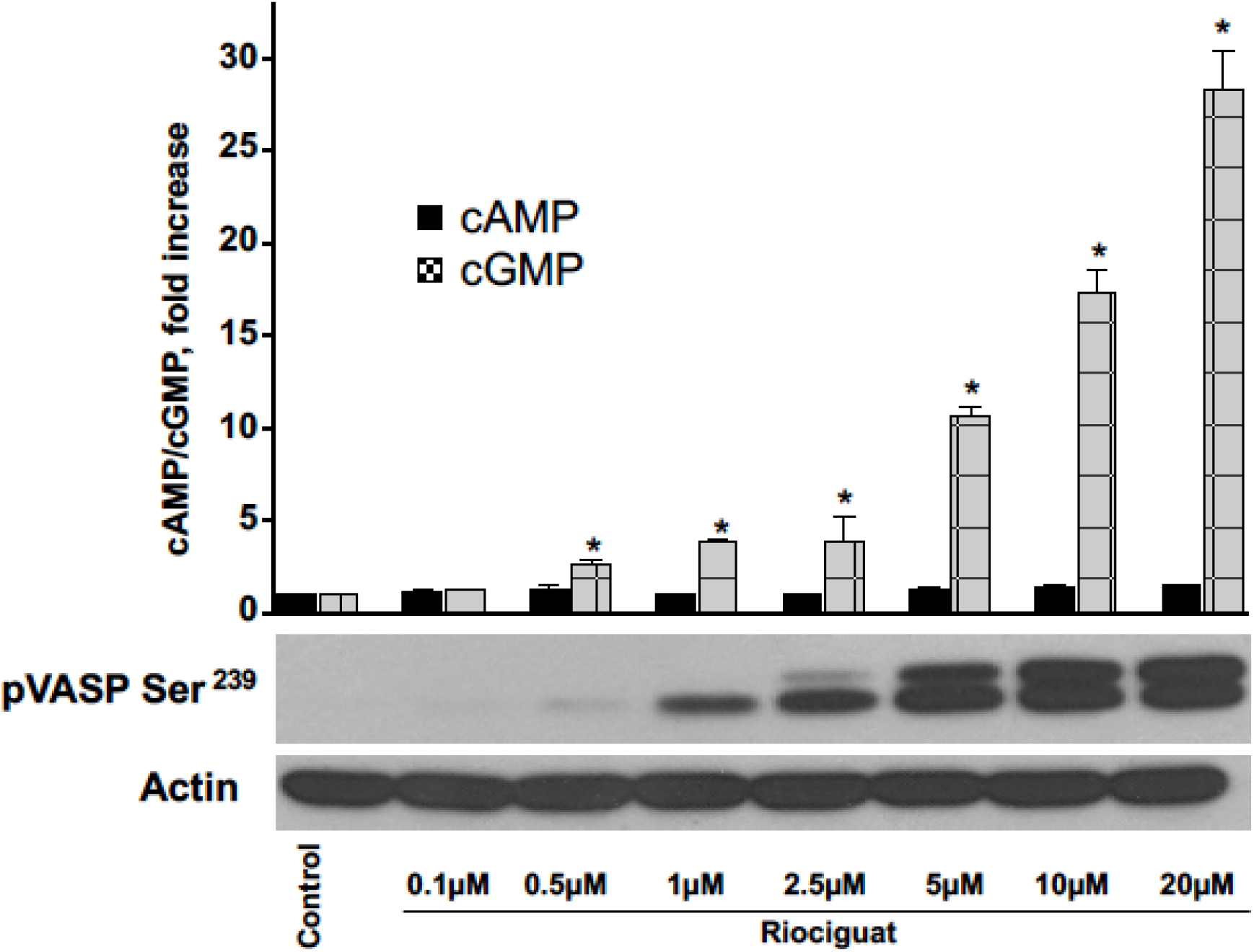
sGC activator Riociguat concentration-dependently increases cGMP without significant changes in cAMP levels in human platelets. Washed human platelets (3×10^8^/ml) were stimulated with indicated concentrations of Riociguat for 5 min. sGC activation was monitored by the analysis of VASP phosphorylation (pVASP^Ser239^ Western blot, actin blot served as a loading control) and cAMP, respectively,cGMP levels by EIA. Results of three independent experiments are presented as mean ± SEM. * p < 0.05 compared to control taken as 1. Shown is a representative blot from three independent experiments.

We observed a significant increase on cGMP levels with increasing concentrations of Riociguat, starting at 0.5 µM while no changes were observed on cAMP levels.. Phosporylation of VASP mirrored this increase on cGMP concentration.

Similarly, increasing stimulation time of sGC with 1 µM Riociguat shows significant increase in cGMP levels as well as VASP phosphorylation compared to control (Figure 7).

**Figure 7.**
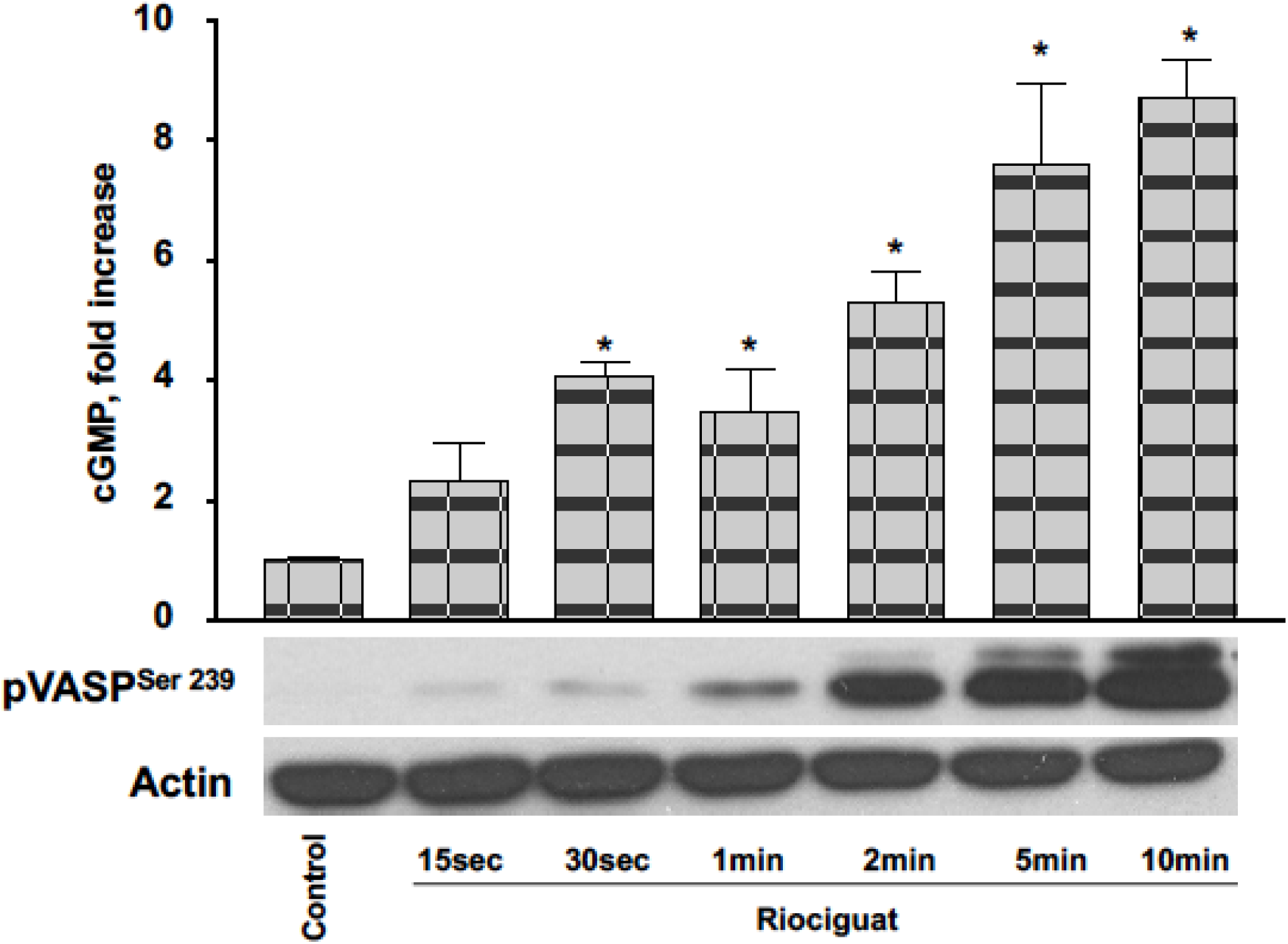
Activation of sGC via Riociguat leads to time-dependent increases of cGMP level and VASP phosphorylation in human platelets. Washed human platelets (3×10^8^/ml) were stimulated with 1µM Riociguat for indicated times. sGC activation was monitored by the analysis of VASP phosphorylation (pVASP^Ser239^ Western Blot, actin blot served as a loading control) and cGMP (EIA). Results of three independent experiments are presented as mean ± SEM. * p < 0.05 compared to control taken as 1. Shown is a representative blot from three independent experiments.

SNP is a NO-releasing compound, thus it reflects activation of sGC pathway on a physiological background. We observed a concentration-dependent (1 nM - 10 µM) significant increase in cGMP levels and VASP phosphorylation, starting with 10 nM SNP (Figure 9). Similarly, a prolonged stimulation time at low SNP concentration (1 µM) showed increased cGMP levels and VASP phosphorylation, starting after 30 sec. (Figure 10).

**Figure 8.**
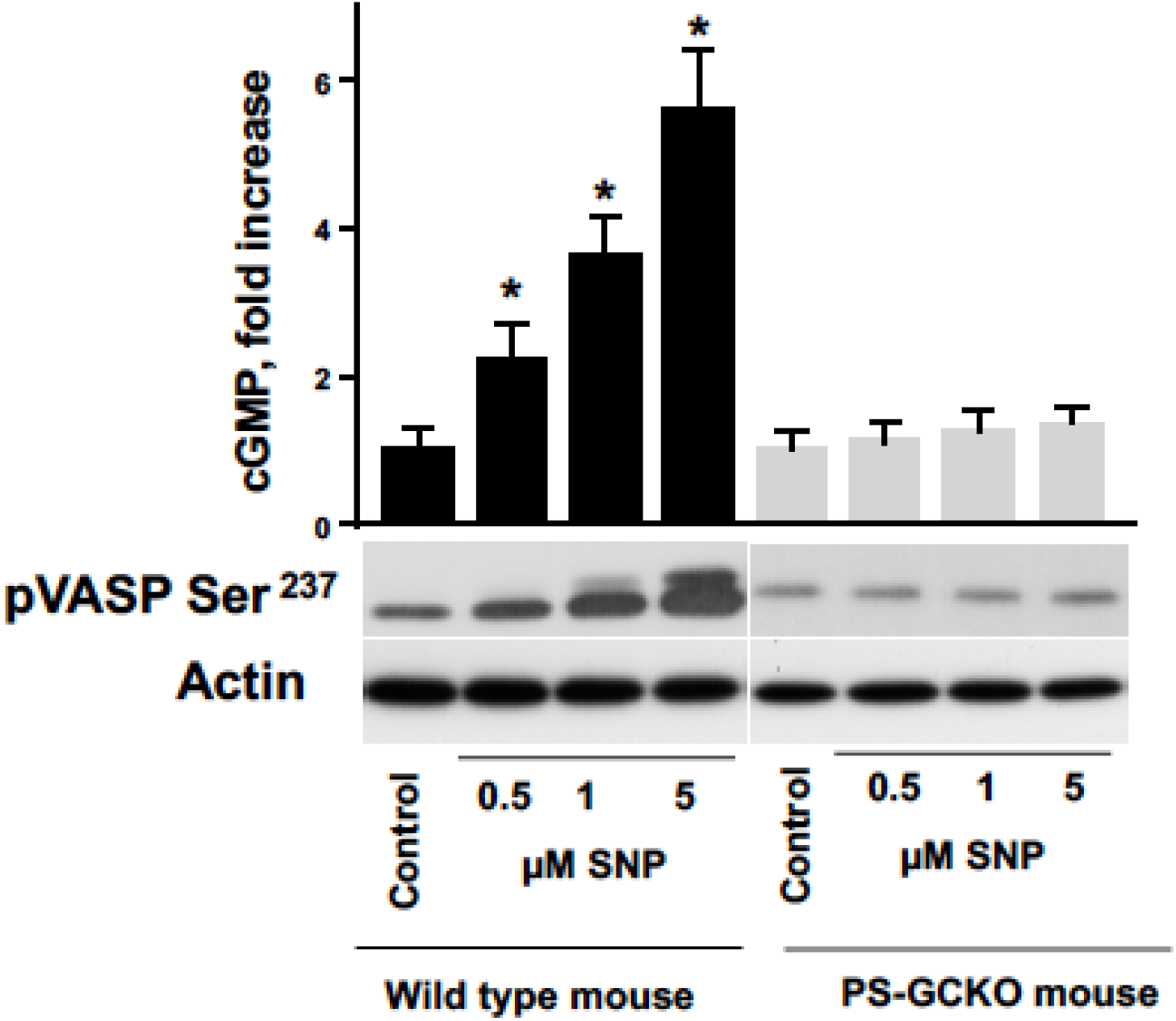
NO donor, SNP concentration-dependently increases cGMP level in Wild type, but not in platelet-specific sGCβ1 knockout (PS-GCKO) mouse platelets. Washed murine platelets (3×10^8^/ml) were stimulated with indicated concentrations of SNP for 2min. sGC activation was monitored by the analysis of VASP phosphorylation (pVASP ^Ser237^ Western Blot, actin blot served as a loading control) and cGMP level by EIA. Shown is a representative blot from three experiments. Results of three independent experiments are presented as mean ± SEM, * p < 0.05 compared to control taken as 1.

**Figure 9.**
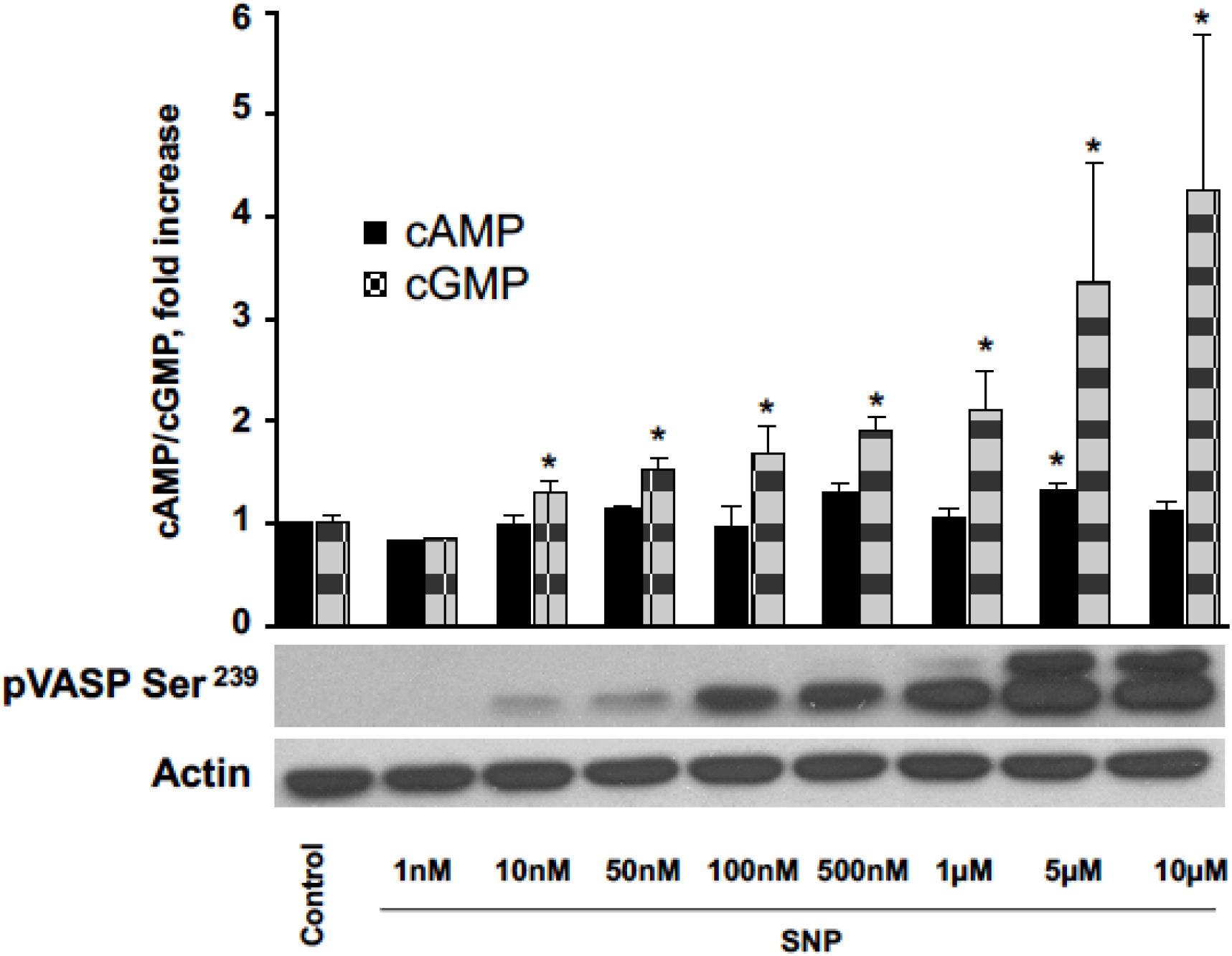
SNP concentration-dependently increases VASP phosphorylation and cGMP level. Washed human platelets (3×10^8^/ml) were stimulated with indicated concentrations of SNP for 2min. sGC activation was monitored by the analysis of VASP phosphorylation (pVASP^Ser239^ Western Blot, actin blot served as a loading control) and cAMP, respectively,cGMP levels by EIA. Shown is a representative blot from three experiments. Results of four representative experiments are presented as mean ± SEM, * p < 0.05 compared to control taken as 1.

**Figure 10.**
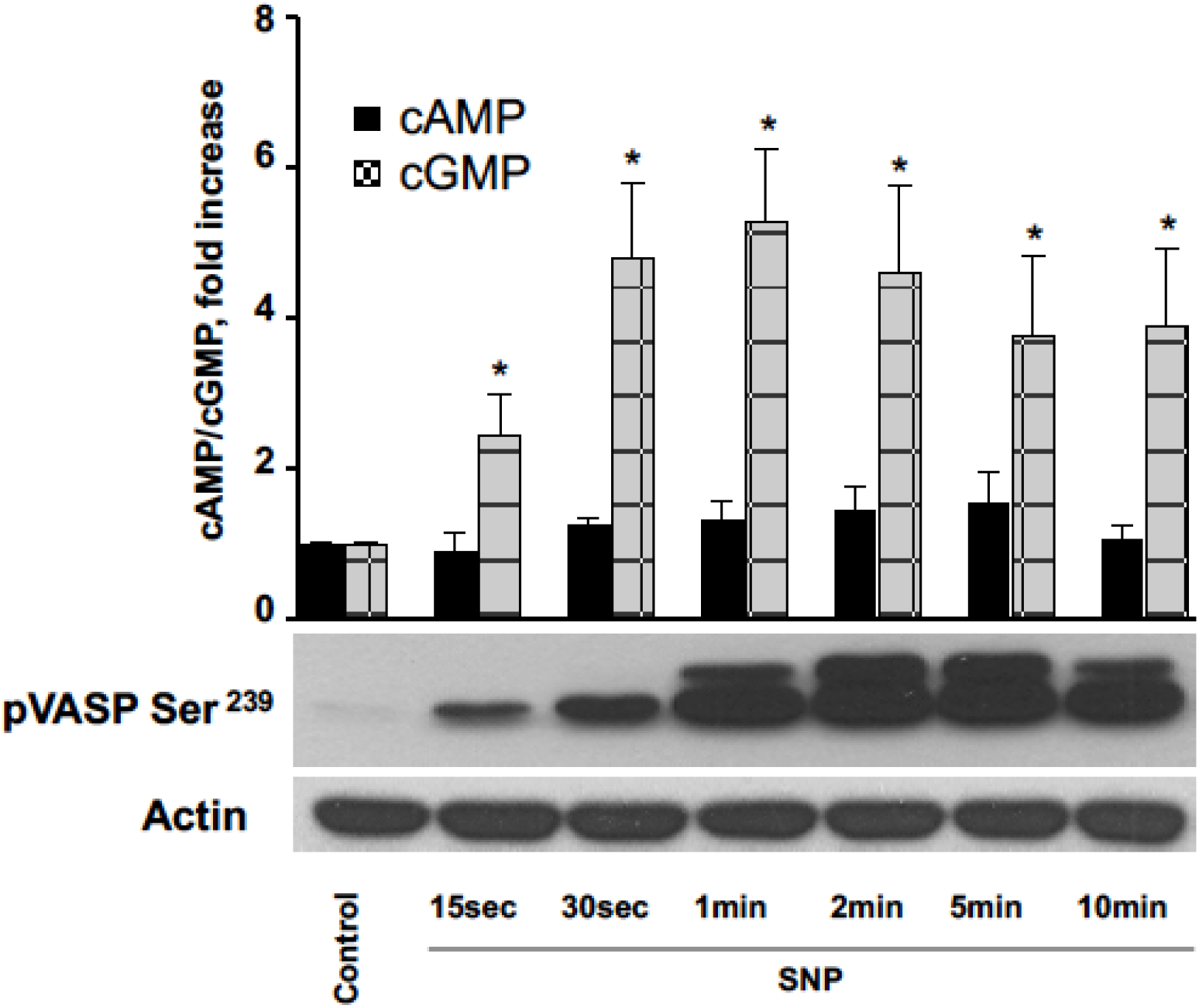
SNP time-dependently increases VASP phosphorylation and cGMP without significant changes in cAMP levels in human platelets. Washed human platelets (3×10^8^/ml) were stimulated with 1µM SNP for indicated times. sGC activation was monitored by the analysis of VASP phosphorylation (pVASP^Ser239^ Western Blot, actin blot served as a loading control) and cAMP, respectively,cGMP levels by EIA. Results of 6 representative experiments are presented as mean ± SEM, * p < 0.05 compared to control taken as 1.

Mouse-models allow to probe here the cGMP levels in more detail by specific enzyme knockouts. Thus, in PS-GCKO mice, cGMP levels and VASP phosphorylation do not change in response to SNP stimulation supporting that the increasing levels observed for cGMP and VASP phosphorylation are depending on sGC activation by SNP (Figure 8).

Finally, also concentration-dependent activation of sGC with Bay 41-2722 revealed enhanced cGMP levels and VASP phosphorylation (Figure 11). Lasting of Bay 41-2722 stimulation, again showed a proportional increase in cGMP and pVASP levels (Figure 12). Overall, VASP phosphorylation occurs later than cGMP indicating that the phosphorylation of VASP needs more time in at least one step of its phosphorylation cascade.

**Figure 11.**
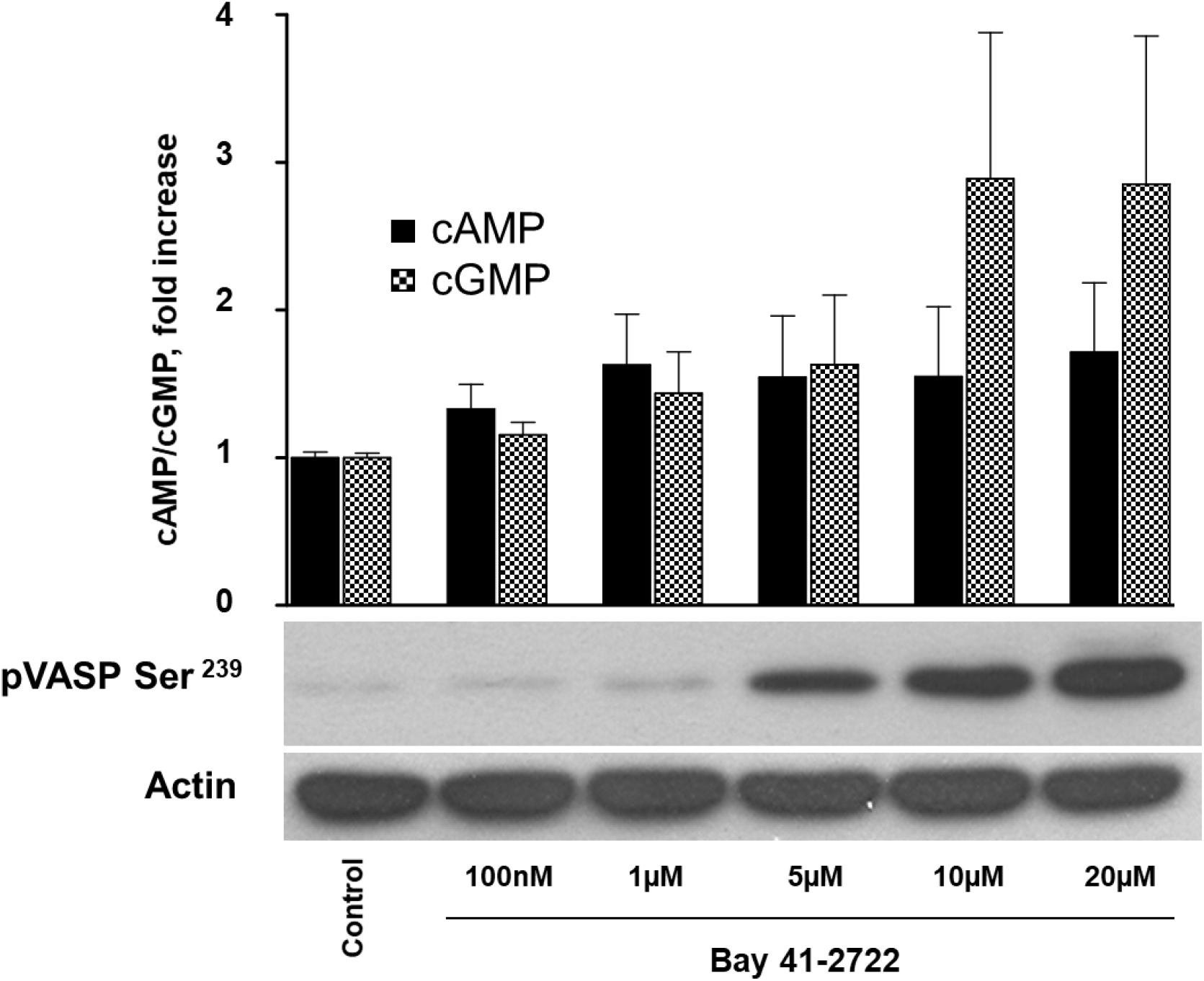
sGC activator Bay 41-2272 concentration-dependently increases VASP phosphorylation and cGMP without significant changes in cAMP levels in human. Washed human platelets (3×10^8^/ml) were stimulated with indicated concentrations of Bay 41-2272 for 5min. sGC activation was monitored by the analysis of VASP phosphorylation (pVASP^Ser239^ Western Blot actin blot served as a loading control) and cAMP, respectively,cGMP levels by EIA. Results of four representative experiments are presented as mean ± SEM, * p < 0.05 compared to control taken as 1.

**Figure 12.**
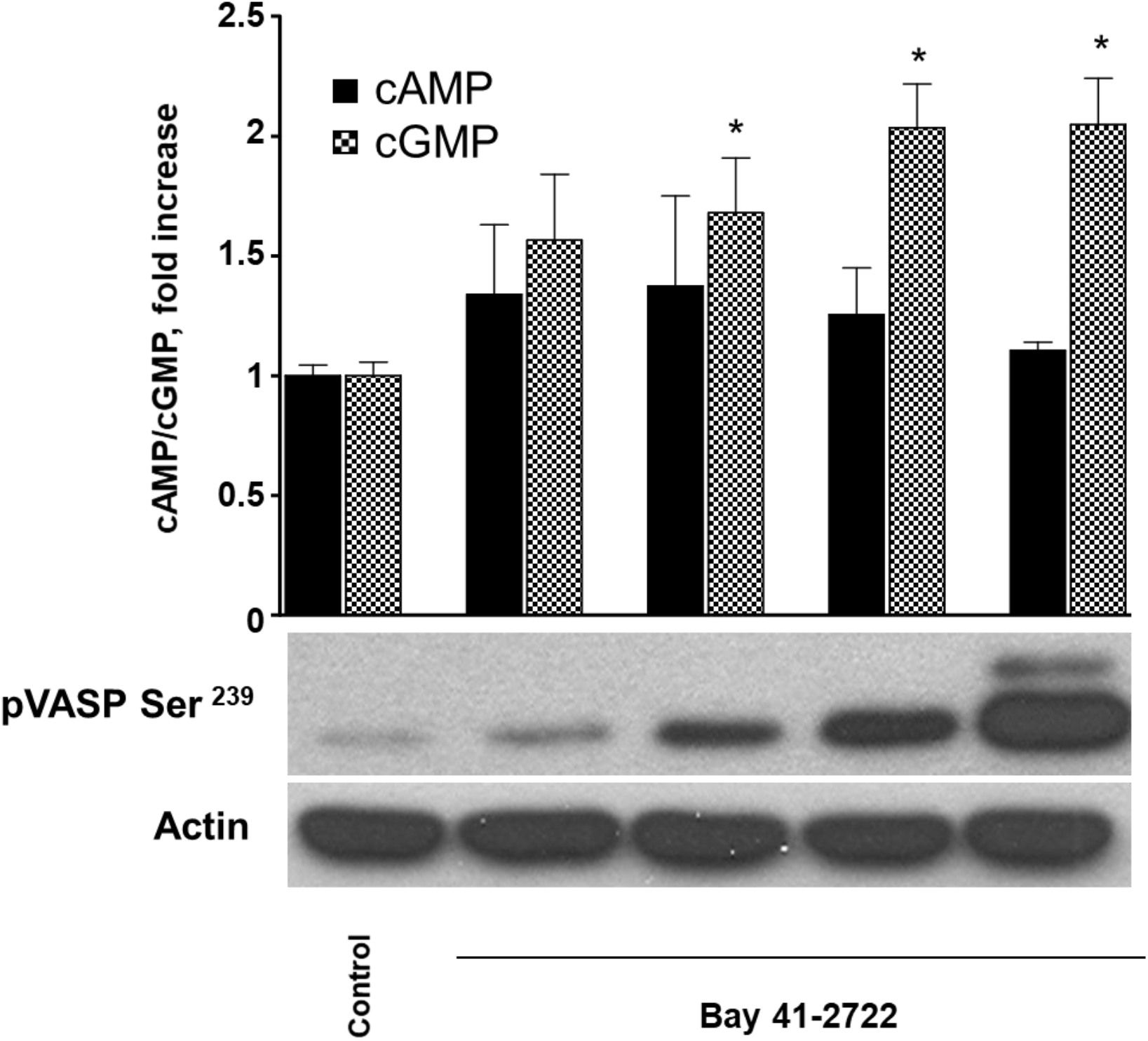
sGC stimulator Bay 41-2272 time-dependently increases VASP phosphorylation and cGMP. Washed human platelets (3×10^8^/ml) were stimulated with 5µM Bay 41-2272 for indicated times. sGC activation was monitored by the analysis of VASP phosphorylation (pVASP Ser^239^ Western Blot actin blot served as a loading control) and cAMP, respectively,cGMP levels by EIA. Results are representative of 4 independent experiments as mean ± SEM, * p < 0.05 compared to control taken as 1.

### Model validation using experimental cGMP time series data

For model validation, we used all the data given as supplemental files (raw_data_Bay_41-2272.xls, raw_data_Riociguat.xls, raw_data_SNP.xls, p-VASP_data.xlsx). Each experiment is performed several times. In each instance of an experiment, the cGMP-levels as well as the cAMP- and VASP-levels if available are measured after several predefined time intervals. From all measurements of an experiment for the same time points, we calculate the mean value, which serves as the best value for the series of measurements, as well as the standard error, which serves as the error bar for the corresponding best value. The curve consisting of best values with the corresponding error bars serves as the time curve for our model validation. We selected the data points for the file data_cGMP.xls whose error bars are not too big compared to the best value and not too small if there are only few measurements. For example, in our opinion it is not reasonable if there are two measurements both (maybe coincidently) with the same value (even though with reasonable values). For the VASP data in the file data_cGMP_VASP.xls, the procedure is the same as described for the cGMP data. However in the VASP case, since the standard error is very inhomogeneous between different stimuli, although the best values are comparable in orders of magnitude, we choose to take the standard error model of Potterswheel (click Settings in the Potterswheel GUI on the left hand-side, the crossed hammer and wrench, then the slider “Fields” and then “data”. There set in the field “yStdCalculations” the value “1 error model of observation” instead of “2 std given in file”) in order not to bias the process of fitting. In the standard error model, the value of the standard deviation equals 10% of the best value of the considered data point plus 0.5% of the maximum value of the observation variable (to which the data point belongs) that the observation variable takes in the data set. We can set the error model in the PW settings as above where we click “model” instead of “data”. Making the error homogeneous prevents overweighting time curves in the process of fitting where the standard error is consistently smaller than with others. This is reasonable since all the data is collected with the same process of measurement and thus each data point of comparable value is supposed to have a similar error. This would be certainly the case if we repeat the measurements more than just three times. However, for the purpose of this work in order to show also principles of modeling, in particular a systematic design of input-output-interfaces, the measurement provides sufficient data.

The data for the time series up to ten minutes for different drugs at a fixed concentration can be seen in Figures 5-7. The time series for different drugs up to two minutes for different concentrations can be seen in Figure 8-10.

In the next step, we present our results from the fitting process with Potterswheel (Maiwald, 2008). Details for the procedure are given in the Methods-Section. In Table 4 we can see the best fitted parameters from the model depicted in Table 3. In Figure 2 we can see the corresponding time curves of the cGMP concentration for different drugs (SNP, Bay, Rio) and concentrations of the drugs simulated with the model from

**Table 4:** Fitted parameters where the x-values are the initial values for the dynamic variables and the k-values the model parameters.

|  |  |  |  |  |  |
| --- | --- | --- | --- | --- | --- |
| $x_2$ | 1 | $k_5$ | 460 | $k_{15}$ | 2.29 |
| $x_3$ | 0.14 | $k_6$ | 0.0006 | $k_{16}$ | 0.55 |
| $x_4$ | 0.22 | $k_7$ | 30 | $k_{17}$ | 110 |
| $x_5$ | 0.08 | $k_8$ | 127 | $k_{18}$ | 1.25 |
| $x_6$ | 0.0005 | $k_9$ | 0.34 | $k_{20}$ | 0.04 |
| $x_7$ | 179 | $k_{10}$ | 187 | $k_{21}$ | 0.001 |
| $x_8$ | 1.25 | $k_{12}$ | 52 | $k_{22}$ | 0.11 |
| $x_{11}$ | 412 | $k_{13}$ | 0.44 | | |
| $x_{12}$ | 16 | $k_{14}$ | 0.0008 | | |

Table 4 with the parameters from Table 4. In addition, in the same color, we have the corresponding data points of the cGMP concentration with its error bars. They are listed in the file data_cGMP.xls in the Supplement. Figure 2 is a screenshot from Potterswheel and is intended to be illustration. As a result, we have that based on the data our model cannot be rejected (subsequently shown) and thus the model, in particular the topology of the interaction network, with the fitted parameters can be used as an explanation for the experiment investigating the connection between cGMP turnover and drug concentration for platelets. Furthermore it is reasonable to use the model for further *in silico* investigations, for example with the optimization framework presented in Breitenbach et al. (2019a,b) for optimal dosage calculations or for finding most effective drug combinations *in silico*.

According to the Potterswheel online documentation (https://potterswheel.de/Documents/PottersWheel-Introduction.pdf), section “Goodness-of-fit”, we see that on a level of significance of 5%, we cannot reject our model for the following reason. There are some values displayed after each sweep of parameter optimization in the MATLAB Command Window. In our case, we have N=36 (number of fitted data points), p=25 (number of fitted parameters), the Chisq=15.2 (Chi-square value), p-value(N)=1, p-value(N-p)=0.17. Since the p-value(N-p) is > 0.05, we can conclude that the model is fitting the data well (as the good fit hypothesis cannot be rejected, see the documentation)). The model can easily be used for further calculations and investigations. Furthermore, it shows that our framework of modeling and including the effect of drugs as external stimuli is useful and functional.

In Table 4 we present the values for the initial values of the dynamic variables and the values for the parameters reasonably rounded. The detailed numbers can be found in the model_cGMP.m file, see the Supplement.

We remark that the coupling constants *k*_20_, *k*_21_, *k*_22_ (coupling between GC and the corresponding drug) differ each in an order of magnitude. The strongest effect is seen with Rio while the lowest effect is observed Bay. We can interpret this according to our mathematical model in a way that the different drugs differ in their effect on stimulating GC. However, this has to be discussed in more detail. Higher activation of sGC by a stimulus is not necessarily connected to a higher affinity and binding constant but can also be explained by a better cell permeability of the drug. This will be discussed in the subsequent paragraph. We would like to remark that also different models are possible to test and to distinguish where the different effects of the drugs come from. For example, we can consider a model that also includes the transport of the drugs into the platelets, maybe an active transport or via diffusion. However, to test different models with respect to such effects and to make statistical significant statements about the validity of models, more detailed data will be necessary. For example data about the concentration of the drug inside and outside a platelet, in the best case time resolved, with the corresponding time-resolved curves of cGMP. In our case, we would like to remark that in order to describe our data, we would like to choose a model with a number of new parameters as small as possible to explain the data and demonstrate that our external stimuli framework is reasonable. Since our model cannot be rejected, it is not necessary to add a more complex reaction mechanism with more free parameters to end up with a more complex model with the same result not being rejected.

The time curves in Figure 2 can be interpreted as follows. At small concentration of drugs GC is stimulated, the concentration of cGMP increases and converges to its equilibrium. At the equilibrium, the basal afflux of cGMP plus the additional afflux by the stimulated GC equals the rate by which cGMP is linearized (hydrolyzed) to GMP by PDE2, PDE3 and PDE5.

If the concentration of a drug reaches a threshold (green curve, Rio at 20 µM), we can see inertial effects. The activation of the PDE2, PDE3 and PDE5 needs some time until there is sufficient active PDE for the linearization of cGMP. Since the additional afflux is bounded by the concentration of GC and thus of GC-stimulated, the linearization of cGMP to GMP by PDEs catches up and thus we have a decay of the cGMP concentration such that the concentration of cGMP also in this case converges to its equilibrium. In the equilibrium, the afflux of cGMP and the linearization to GMP by PDE equals each other. The peaks for high concentration of drugs are also seen in further experimental studies (Mullerhausen a,b). Furthermore, we remark that the software Potterswheel fits the total model to the given data points considering the corresponding standard error bars. Since the parameters of the model are fit such that the chi-square value is minimized and make an optimal fit, it is not necessary that all data points are on a curve, see Fig. 2, light green curve. This can in particular happen if the error bars are big compared to those of other data points indicating that the data point with the big standard error is not as reliable as the ones with small error bars. To evaluate the model in total the p-value is a measure that links the chi-square values with the number of free parameters and provides information if the model has to be rejected (as too many data points are too far from their predicted position).

Since our model is based on the core model of Wangorsch et al. (2011), we additionally validated the topology of the cGMP part of the model by our core experimental data while considering cAMP- and VASP-levels for other parts of the model.

We remark that comparing reaction constants from a model with the same topology, but fitted to data collected from different experiments, although from the same biological system, e.g. platelets, is delicate since the reaction constants depend on factors that may differ from each experimental setup. For example, temperature of the setup, the way the platelets have been prepared and can be taken from different animals. However, the topology, that means the basic reaction mechanisms (kind of enzymes, reaction chains) are the same for all the setups up to constants that scale the curves to the specific data, which means to the differences in the experimental environment. This is possible as far as the topology of the model captures the essential dynamic of the biological system according to which the considered biological system evolves.

The importance of the topology of a model with just constants to be fitted, is very plausible considering for example patient specific therapies where various living conditions or different gene pools can cause different reaction parameters. As a result, the same enzyme has different turnovers for a certain substrate maybe due to a disease, aberration or different alleles of a gen. But for all these cases it is reasonable to use the same topology to describe the corresponding data because e.g. protein cascades are the same. This example stresses that the model topology is crucial to capture the main dynamic of a biological system up to constants that can be fitted to a specific situation. On the other hand, this illustrates that one has to be really aware of the effects that influence reaction constants of a model. Moreover, one has to make sure just to use data collected in a comparable environment for fitting each model if the purpose of the modeling is to compare the reaction parameters of models, e.g. same temperature or the cell suspension consists of the same ingredients. In our work, the purpose is to demonstrate a systematic process for modeling external stimuli and thus providing a rational way of creating model topologies to include external stimuli into modeling. In addition, we will show below how to combine different models module wise to describe a system with well validated sub models.

The considerations above hold for the case whether a model is sensitive or not sensitive with respect to a certain reaction parameter. For the parameters to which a model is not sensitive, we additionally have the effect that we can have (almost) the same simulated curves for different parameters. The notion of this work is that we need only one model describing our data such that it cannot be rejected for further *in silico* studies. The exact value of certain reaction parameters is not critical as long as the total model cannot be rejected. The reliability of the predictions and results of simulations increases with the quality of the data base, which means to have many data points with only little error bars collected from experiments with comparable conditions. The number of models that cannot be rejected decreases with the increasing quality of the data. Consequently, we also need to refine models to capture also the mechanisms with small effects encoded and only resolved in high quality data where they can be distinguished from white noise. Thus, our proposed procedure for mapping input information of a biological system to output information provides more reliable input-output-relations the better the data is.

Reliability in our context means that the probability that a model with certain reaction constants fitted from a set of high-quality data can also not be rejected based on a different set of data collected from the same experimental setup with comparable conditions where the data for the fitting process has been collected.

If there is a sufficient amount of data, we can split the data into a training set and a test data set. On the training data set we fit the parameter such that the corresponding model cannot be rejected. On the test data set we just evaluate the fit statistics, e.g. with Potterswheel, and see if the model with the parameters fitted on the training data and now fixed cannot be rejected based on the test data set either.

Next, we present the results from taking the cGMP time curves as an output of the model depicted in Table 4 and input it into the models depicted in Table 2 and Table 3. For this purpose, we fix the initial values of the dynamic variables and the parameters that are shown in Table 4 by changing the fitSettings in the model file “model_VASP.m” and “model_VASP_wangorsch.m” from “global” to “fix”. Consequently, only the new model parameters are fitted by Potterswheel. In this way, we demonstrate how to use a validated model, here the cGMP model, with the cGMP concentration *x*_2_ being the output variable and concomitant, serving as an input variable for another model that is fitted based on the validated model.

The fitting process is analogous to the model of cGMP. As a result we have for the model “model_VASP.m” a p-value of 0.77 and a p-value(N-p)=0.51. Also for the model “model_VASP_wangorsch.m”, we obtain a p-value=0.64 and a p-value(N-p)=0.12. This indicates that both models cannot be rejected based on the data, they are a statistically well confirmed fit to the data, and this is confirmed at a level of significance of 5%. This demonstrates that our framework of combining different models via an input-output-interface works and this example validates it. For illustration, we depict the corresponding time curves in Figure 3 and Figure 4 that are a screenshot of Potterswheel. The exact values of the corresponding model parameters can be found in the model files “model_VASP.m” and “model_VASP_wangorsch.m”.

We remark that in our data, the cAMP concentration was basically constant. As already implemented and apparent from Wangorsch et al. (2011) the corresponding PKA is activated and provides a constant afflux of phosphorylated VASP. We tested a further model by adding a free parameter to the equation modeling the evolution of *x*_15_ (phosphorylated VASP) and subtracted it from the equation modeling the evolution of *x*_14_ (unphosphorylated VASP) in the model depicted in Table 2. We cannot reject this model either but since we cannot reject models that do not explicitly consider this constant afflux of phosphorylated VASP from the cAMP pathway, we can reasonably neglected this constant afflux here in our models. Furthermore, we could also not reject the fitted model from Table 2 where we exchanged the terms modeling a Michalis-Menten dynamics by the corresponding terms based on the law of mass action. By this step, we just take the enumerator and replace the corresponding denominator by 1. Consequently, we see that we cannot reject the model based on the law of mass action and do not need to make any assumptions that are used to model Michaelis-Menten-dynamics. This is also in agreement with the fact that the basal model depicted in Table 3 cannot be rejected where we have just bilinear terms considering that a reaction rate is proportional to the involved reaction partners. The fact that we can work just with two equations can be explained by the fact, that the concentration of PKG is quite constant compared to the time scale on which we consider the process and thus is considered in the free reaction parameter *k*_24_.

## Discussion

We present here an overarching, unified model of cyclic nucleotide pathways with a special focus on cGMP effects and that is validated by direct experimental measurements. For this purpose, it is necessary and sufficient that the model cannot be rejected based on the data available and a statistical level of significance. Once the model is set up and describes the data curves sufficiently well, i.e. the model cannot be rejected, we can further exploit the information coded in the fitted mathematical model by an optimization framework presented in Breitenbach et al. (2019a,b) in order to calculate an optimal drug dosage or most effective drug combinations for a desired cGMP concentration or for steering the concentration time-resolved along a desired cGMP concentration curve.

Furthermore, our model is fit to each drug and by the virtue of the mathematical structure of the model, there is also information about the combination effects of drugs included. Consequently, we can search fully automatically in silico for the most effective drug combination in connection with the optimization framework mentioned above and save a lot of experimental effort. This effort would arise from making experiments for any combination instead of just testing the calculated combination. Finally, we show how to build a comprehensive model from different sub-model modules that in consequence also describes VASP phosphorylation where the cGMP concentration is considered as an input for the VASP model.

We remark that the module wise combination of sub-models is not limited to systems of ordinary differential equations. Also machine learning models that map input to output information can be combined with other machine learning models or systems of ordinary differential equations by the corresponding input-output interface. In order to use such models for calculating optimal external stimuli as in Breitenbach T. et al. (2019a,b), the gradient of these models with respect to the input variables (external stimuli) is needed. Since the explice functions of machine learning models are not always given, the calculation of the gradient is hindered. However, since the model is formulated as a computer program which can be seen as a function mapping input to output signals, the technique of automatic differentiation of programms can be applied providing the required gradient also in the machine learning case. A computer program consists of the concatenation of elementary operations and functions, and if they are all differentiable, by the chain rule any software program and thus any model encoded by such a program is automatically differentialbe. Consequntly, any optimizaiton framework can be applied. For an automatic differentiation framework, see for example Python Tensorflow (automatic differentiation).

So far, we have created our model based on ordinary differential equations and the available experimental data given in this paper. On a level of significance of 5%, we cannot reject the model. However, this is supposed to enforce us to coordinate experiments and mathematical modeling even more. On the one hand, one needs to provide better data with a smaller standard error and on the other hand, based on less variant data, maybe falsify simple models (have to be rejected based on the p-value) with only a little number of free parameters and simple reaction mechanisms and test more accurate and more complex models.

Next, we discuss our choice of the model regarding how to include drugs. We choose a law of mass action mechanism to describe the interaction of the drugs with the enzyme GC. An alternative would be to choose a Michaelis-Menten reaction mechanism. However, we see advantages to model with the law of mass action for the following reasons:

Firstly, for one Michaelis-Menten term, we need always two free parameters per drug. One is the maximum reaction rate and the other one is the Michaelis constant. Consequently, in our case, we save one free parameter. Secondly, to derive the Michaels-Menten-terms from substrate-enzyme-reactions, we need to make further assumptions from which we do not know if they are fulfilled. The law of mass action is basal to the Michaelis-Menten-theory, that’s why we operate with less assumptions in our modeling.

Once the GC is stimulated, this response does not differ by which drug it was stimulated. This could be done by replacing the equation of GC-stimulated in Table 1 by an equation for each species and add three different terms in the equation for cGMP instead of *x*_12_ multiplied each with another parameter to adapt the effect of each drug-GC-complex. However, since our simpler model cannot be rejected, we do not need this differentiation based on the data for the purpose of the paper to show how to systematically map input information from concentrations of drugs to output quantities.

Next, we discuss more model extensions for covering more cases in the following.

## Model extensions

### Inhibiting effects

We discuss how to extend our model to also include drugs that inhibit the GC. To the model from Table 1, we add one equation for the quantity modeling the species of GC inhibited, e.g. *x*_13_. Then we can replace *k*_12_ by *x*_11_, multiply *x*_12_ by another parameter to adapt for the higher cGMP turnover of GC-stimulated. Analogously, we add *x*_13_ multiplied by a free parameter to fit to the increased turnover of the species GC-inhibited.

However, since we only have the stimulated species, we decided to keep *k*_12_ and just add a term *x*_12_ in the equation for cGMP (*x*_2_) to safe another parameter to obtain a model, which cannot be rejected, with a number of parameters not bigger than necessary.

If there are several drugs, which stimulate or inhibit GC to different degrees we have to replace the equations for GC-stimulated and GC-inhibited by an equation for each species depending on the drug. Then add the corresponding dynamic variable multiplied with a further parameter to the equation for cGMP.

Another possibility to include the inhibiting drugs is to subtract *x*_13_ (species of inhibited GC) from the equation for *x*_2_. The idea is that the cGMP afflux of GC modeled by *k*_12_ decreases. However, the disadvantage of this approach is that we have to care during the fitting process that *k*_12_ ≥ *x*_13_ since the GC does at least produce no cGMP but never lowers the cGMP actively in our model which would correspond to *k*_12_<*x*_13_.

### Drugs with modeled dynamic and building blocks of models

In the next step, we discuss how to include the case in our framework if drugs have their own dynamics, like transport into cells, invasion into an organism via different organs, excretion, conversion or are metabolized in an organism. In this case, we have to model the concentration of the drug as a dynamic variable with an ODE that models its dynamic. In the equations presented in Table 1 the external stimuli *u*_1_, *u*_2_, *u*_3_ are then replaced by the corresponding dynamic variables where the external stimuli are now in the equations modeling the dynamic of the drugs and model the administration of the drugs. For example, the dosage that is applied to the organism. For instance, the administration could be modeled as a pulse of an external stimulus (constant function for a fixed interval of time) corresponding to the dosage. The parameters in the model system describing the dissolution and the distribution of the drug into the organism, in particular these parameters multiplied with the external stimuli functions, can be fit to the data such that the pulse is calibrated. We see that we can work here module wise. Since for example the model, which is fit to the data how much drug is in the blood after administration, can be used for further models describing the metabolism in the liver or for models describing the effect of the drug in an organ. Here the concentration of the drug in the blood is then an input function for the other models. In each model, the external stimuli model the input information into the corresponding model. This means, when working in this module wise manner, the external stimuli can either be directly under control of the experimenter, for example if they have methods to actively add or remove a drug, or indirect via other models. These models represent the possibilities of the experimenter’s influence where the functions of the external stimuli are replaced by the corresponding output functions of that model in which the experimenter can directly control quantities.

By this procedure, we have a building block principle where different models are coupled by replacing the external stimuli functions from one model by the corresponding output functions of a different model modeling the dynamics of the corresponding quantity. This quantity is an external stimulus for the former model. This also has the advantage, that if a model is well validated, for example the uptake of a drug, we just have to fit the parameters for the later model and check if the combined model cannot be rejected. If it is rejected, then we can check each sub-model and see if it can be rejected or not by measuring each external stimulus for each sub-model and input these functions into each sub model calculation.

Another application of this building block framework is the cell-cell interaction where the cells can be seen as sub systems modeled with well validated sub models for each cell. The signal proteins of one cell is on the one hand the output signal of this cell and on the other hand, serves as the input signal for a different cell or cell type. If we consider a single cell or a cell type depends on the model. One cell type can have different expression patterns, for example an inhomogeneous tumor. To describe this, we can use different models each for such a cell with a different mutation leading to a significant different expression pattern. Analogously, we can model different cell types each with a different model, like T-cells communicating with B-cells and in this way build a model of the immune system from the sub systems, which are the different cell types. By this procedure, we can model the immune system based on the different immune cell types and validate each sub model and subsequently the total model of the immune system.

This also makes the identification of potentials for improvement of models much more efficient since we can break down the analysis just to that small part of the model that did not pass our tests. This procedure keeps the investigation clear and rational. Consequently, data can be collected needs-based just where further information for model improvement is needed. The well validated models can then be recycled for further projects.

An application and example of the concept discussed above is that once we have a model for the drugs describing the uptake of the drug into the blood, we can combine this model with our presented one, where we replace the external stimuli functions by the corresponding concentration of the drugs in the blood. Consequently, we create models to perform *in silico* studies about optimal therapies for diseases caused by a misbehavior of the platelets.

The demonstration of the building block procedure was performed by combining the cGMP model with the VASP phosphorylation models.

### Building more detailed models

In this paragraph, we discuss how to model more accurately in the case if the bilinear structure of external stimulus multiplied by a dynamic variable is not sufficient. We remark in our presented case, this is not necessary since the bilinear model cannot be rejected based on our data. To test more accurate models in future work, more accurate data is needed. That means the number of repetitions of a single experiment must be increased to lower the error bars or use techniques of measurements that allow measurements that are more precise.

In case it is necessary to model, for example, cooperativity effects and still keep the bilinear structure we can proceed as follows. For each molecule of drug that binds to an enzyme GC, we create a new species that again reacts with a further molecule of the same drug or (also possible) with a different drug according to the law of mass action. For each intermediate species, we have one equation with the bilinear structure consisting of a product of the enzyme complex with the already bound molecules and the external stimulus function (e.g. modeling the concentration of the molecule) to be considered where the bilinear reaction mechanism according to the law of mass action is applied. Thus, also more complex reaction models can be traced back to the presented bilinear model structure analogous for GC and GC-stimulated, see Table 1. In this way, cooperativity effects can be modeled. However, such detailed models of reaction mechanisms are only necessary if models with compressed mathematical reaction mechanism have to be rejected based on the available data where a solution may be to go to more detailed models with more parameters. By the procedure of breaking down complex models to the bilinear approach, the model can then be directly combined with other frameworks like the optimization framework presented in Breitenbach T. et al. (2019a,b) to optimize therapies, to optimize searches for effective active ingredients and their dosage.

## Conclusions

In this work, we focused on developing a model describing the time curves of measured cGMP levels in platelets. The information coded in the mathematical model can be used for further *in silico* studies like optimal dose calculations or most effective drug combinations. For this purpose, it is not necessary to perform sensitivity analysis of the model to evaluate the ranges for the model parameter for further studies with them, but only need to describe the time curves with a model that cannot be rejected.

We extended the model from Wangorsch et al. (2011) as well as other more recent models (Kleppe et al., 2018) with our external stimuli framwork. This allows to model drug effects and to validate the total model with our experimental data. Based on our experimental data the model is confirmed. Moreover, the model is even further validated by applying its output into a VASP phosphorylation model, which also is confirmed according to our corresponding VASP phosphorylation data.

This work is supposed to encourage more specific and accurate measurements on nucleotide and cGMP effects in platelets as well as to measure the values of more dynamic variables or quantities to create more detailed models and thus refine our imagination of the investigated biological processes. Furthermore, coordinating experiments with modeling leads to purposefully measurements and planning of experiments. A coordinated process of experiments and mathematical modeling results in more accurate models and in turn motivates more specific measurements, which will focus the experimental effort only to collect that data where information is missing for the understanding of the whole investigated biological process. Only collecting specific data can accelerate the gain of insight since experiments can now be done faster and thus become more efficient with our model. Finally, our focus on external stimuli should stimulate at the same time the development of further pharmacological targeting of platelets via the cyclic nucleotide pathways for improved treatment of cardiovascular disease.

## List of abbreviations

cAMP: cyclic adenosine monophosphate
AMP: adenosine monophosphate
cGMP: cyclic guanosine monophosphate
GMP: guanosine monophosphate
AC: adenylyl cyclase
GC: guanylyl cyclase
PDE: phosphodiesterase
PKA: cAMP-dependent protein kinase
PKG: cGMP-dependent protein kinase
VASP: vasodilator stimulated phosphoprotein
GPCR: G-protein-coupled receptor
ODE: ordinary differential equation
sGC: soluble guanylyl cyclase

## Authors’ contributions

TB designed the extension of the basic model, performed the mathematical modeling and the parameter fitting with Potterswheel. TD lead and guided the study.

TB, NE, ÖO, GW, EB, RF, MD, SG and TD were involved in the data analysis.

NR and SG conducted all experiments for the experimental data shown.

NE, EB, RF, AF, SG provided cGMP experimental analysis expertise.

TB, ÖO, GW, MD and TD provided cGMP modeling expertise.

TB, MD, SG and TD were involved in drafting the paper which was followed by editing and comments by all authors. All authors read and approved the final manuscript.

## Acknowledgements

The authors gratefully acknowledge the support by Deutsche Forschungsgemeinschaft (DFG) Project number 3 74031971 – TRR 240 [ÖO:Z2, signalling][RF:A5, cGMP][TD:INF, software]

## Availability of data and materials

All results are contained in the manuscript and its supplementary files including all details on the modeling involved and all experimental data for the study.

## Materials and Methods

### Platelet preparation

#### Chemicals

SNP, sGC activator Bay 41-2272 and sGC inhibitor ODQ were obtained from Sigma-Aldrich (Seelze, Germany). Riociguat (Bay 63-2521) was a kind gift from Bayer (Wuppertal, Germany). Anti-phospho VASP^Ser239^ antibody was from Nanotools (Teningen, Germany), Anti-phospho PDE5^Ser92^ antibody was a kind gift from Dr. S. Rybalkin (Department of Pharmacology, University of Washington, Seattle, USA). Polyclonal rabbit anti-actin antibody was from Santa Cruz Biotechnology (Heidelberg, Germany). Horseradish peroxidase conjugated goat anti-rabbit and anti-mouse antibodies were from Bio-Rad Laboratories, Inc. (Munich, Germany).

#### Human platelet preparation

Human platelets were prepared and used as previously reported (Gambaryan et al, 2004) with small modifications. Blood was obtained from healthy volunteers according to our institutional guidelines and the Declaration of Helsinki. Our studies with human platelets were approved by the local ethics committee of the University of Würzburg (Studies No. 67/92 and 114/04).

Blood was collected into 1/7 volume of ACD solution (12 mM citric acid, 15 mM sodium citrate, 25 mM D-glucose and 2 µM EGTA, final concentrations). Platelet rich plasma (PRP) was obtained by 5 min centrifugation at 330g. To reduce leukocyte contamination, PRP was diluted 1:1 with PBS and centrifuged at 240g for 10 min. Subsequently, the supernatant was centrifuged for 10 min at 430g, then the pelleted platelets were washed once in CGS buffer (120 mM sodium chloride, 12.9 mM trisodium citrate, 10 mM D-glucose, pH 6.5), and resuspended in HEPES buffer (150 mM sodium chloride, 5 mM potassium chloride, 1 mM magnesium chloride, 5 mM D-glucose, 10 mM HEPES, pH 7.4). After 15 min rest in a 37 °C water bath, washed platelets were used for experiments. Washed platelets (WP) were used at a concentration of 3 x 108 platelets/ml for Western blot analysis.

#### Mice

Platelet specific sGC knock out mice (PS-GCKO) were generated as previously described (<u>Rukoyatkina</u>*<u>et al</u>*<u>., 2011</u>). All mice were housed according to the German Animal Welfare Act in a barrier facility with a 12-hour light-dark cycle. Mice were kept in EU Type II IVC cages with a maximum of 5 mice per cage under conventional conditions. Mice were backcrossed on a C57Bl/6 background. The mice used in the experiments were 8-week-old WT and platelet-specific sGC KO male mice.

#### Mouse platelet preparation

**Blood was collected from the orbital sinus of WT, and PS-GCKO mice anesthetized by isoflurane. PRP and WP were prepared according to published protocols (Rukoyatkina et al**., **2011). Briefly, blood was collected into 1/7 volume of ACD and centrifuged at 300g for 5 min. PRP was centrifuged for additional 8 min at 80g to separate from contaminating erythrocytes. Purified PRP was centrifuged at 700g for 5 min. Pelleted platelets were resuspended in Tyrode buffer (137 mM NaCl, 2 mM KCl, 2 mM MgCl2, 12 mM NaHCO3, 0.3 mM NaH2PO4 x2 H2O, 5.5 mM D-glucose, 5 mM Hepes, pH 7.4) and the number of used WP for Western blots experiments was adjusted similar to human platelets**.

#### cAMP and cGMP Measurements

Levels of cAMP and cGMP were evaluated using a cAMP EIA kit and cGMP EIA kit, respectively, following the manufacturers’ instructions (Cayman Chemical, Hamburg, Germany).

#### Western blot analysis

Western blot analysis was performed as described in details (Gambaryan et al, 2010). Briefly, washed platelets were stimulated for indicated time/ concentrations of used compounds and stopped by adding SDS gel loading buffer. For Western blotting, cell lysates were separated by SDS-PAGE, transferred to nitrocellulose membranes and the membranes were incubated with appropriate primary antibodies (phospho-VASP-Ser^239^ phosphoPDE5, and Actin) overnight at 4°C. For visualisation of the signal, goat anti-rabbit or anti-mouse IgG conjugated with horseradish peroxidase were used as secondary antibodies, followed by ECL detection. The blots were analyzed densitometrically using NIH Image J software for uncalibrated optical density.

#### ANOVA analysis of the experimental data

All experiments were performed at least thrice, and data are expressed as means ± SEM. Differences between groups were analyzed by ANOVA, and the Student’s t-test was used when appropriate.

### Systems biological modeling

We used the **MATLAB toolbox Potterswheel** (Maiwald, 2008) to fit our model to the experimental data (data_cGMP.xls, data_cGMP_VASP.xls). For this purpose, we used the fminsearch and the Trustregion method to optimize the parameters. We iterated between these two methods and started each optimization sweep for the parameters with the results from the previous sweep. Between these sweeps we set some parameters manually to reasonable values with the PW Equalizer if necessary. For example if some reaction constants or initial values of the quantities were negative as long as we did not see convergence to reasonable parameters. For the integration of the ordinary differential equation we used the ODE15s method. For the tolerance of the linesearch method we used tolFun=tolX=10^−9^ and for tolFun=tolX=10^−9^ for the Trustregion method. This settings can all be done in the PW Configuration (can be entered for example via the Potterswheel GUI, on the left by clicking on the image with the crossed hammer and wrench. Next click on the slider “Field” in the bar on top of the opening tab).

We provide the **model and data files** to enhance the confirmability and reproductivity of the results. Furthermore, the user can easily modify the files and make own experiments. Once a MATLAB and Potterswheel version is installed, the user can immediately start by entering one after another “pwAddModel(‘model_cGMP.m’)”, then open the PW Configuration in the PW Window, click Fields, then data and choose “2 std given in file. Then go to the MATLAB Command Window and enter “pwAddData(‘data_cGMP.xls’)”, “pwCombine”, “pwArrange” and “pwFit” into the MATLAB Command Window. For the VASP models proceed analogously.

model_cGMP.m file – the model file for Potterswheel for the cGMP model

model_VASP_wangorsch.m file – the model file for Potterswheel for the VASP model

model_VASP.m – the model file for Potterswheel for the basal VASP model

data_cGMP.xls file – the data points for the fitting with Potterswheel for the cGMP model

data_cGMP_VASP.xls file – the data points for the fitting with Potterswheel for the VASP models

raw_data_Bay_41-2272.xls – raw data of cGMP concentration for Bay

raw_data_Riociguat.xls – raw data of cGMP concentration for Rio

raw_data_SNP.xls – raw data of cGMP concentration for SNP

raw_data_p-VASP.xlsx – raw data of p-VASP concentration for Bay, Rio, SNP

### Model fit to the data

The **χ**^**2**^ -value given for each model shown in Fig. 2-4 is a measure for the distance between the model prediction and the corresponding data points. Consequently, the smaller the value is, the more likely it is that the model explains the data. Considering the number of data points (N=36), the p-values can be calculated giving the probability to obtain the value of **χ**^**2**^ assuming that the hypothesis that the model explains the data is true. The p-value gives an estimation if the model based on the data has to be rejected or can be reasonably used for further calculations. In detail we have for Fig. 2: The p-value considering the degrees of freedom (number of data points minus parameters to be fit) is 0.17. Consequently on a level of significance of 5% the hypothesis that the model explains the data cannot be rejected since such a **χ**^**2**^ -value of at least 15.21 with 36-25=11 degrees of freedom appears in up to 17% of the cases given that hypothesis is true. For Fig. 3 the p-value considering the degrees of freedom (number of data points minus parameters to be fit) is 0.51. Again, the hypothesis of a good data fit and good model cannot be rejected. For Fig. 4 the p-value considering the degrees of freedom (number of data points minus parameters to be fit) is 0.12. Again, the hypothesis of a good data fit and good model cannot be rejected.

